# Protozoan Anaerobic Metabolism Emits Acetaldehyde as a Volatile Infochemical Driving Bacterial Biofilm Dispersal

**DOI:** 10.64898/2026.05.25.727473

**Authors:** Prem Anand Murugan, Smruti Mahapatra, Meirav Gefen, Serge Ankri, Ilana Kolodkin-Gal

## Abstract

Predator–prey interactions are classically governed by physical contact or lytic enzymes, leaving the role of small diffusible metabolites as long-range warning cues poorly understood. Because amoeboid protists feed directly on biofilms, assembling a sessile community where predators dominate is ecologically disadvantageous. Here, we show that the mitochondria-lacking parasite *Entamoeba histolytica* emits acetaldehyde via EhADH2, which functions across both liquid and gas phases as a long-range interkingdom warning cue—analogous to bacterial quorum sensing—directly linking a parasite-specific metabolite to microbiome behavior. This infochemical suppresses biofilm assembly and promotes dispersal in *Bacillus subtilis* and related probiotic species. Mechanistically, acetaldehyde selectively rewires cyclic di-AMP (c-di-AMP) homeostasis to relieve SinR/AbrB-mediated motility repression and drive rapid flagellar escape prior to physical encounter, uncovering a conserved second-messenger hub governing cross-kingdom behavioral plasticity

## Introduction

*Entamoeba histolytica* remains a significant global health threat; according to the Global Burden of Disease estimates, Entamoeba-associated infections exceed 2.5 million cases and contribute to approximately 33,000 annual deaths worldwide^1^. As a primary causative agent of amoebic dysentery^2,3^, *E. histolytica* navigates the complex, biofilm-rich environment of the human gastrointestinal tract. In this niche, the parasite exists in constant contact with the gut microbiota- a dense microbial community that serves as a cornerstone of host metabolism, nutrient acquisition, and immune homeostasis ^4,5^. The introduction of *E. histolytica* into this intricate ecosystem via the fecal-oral route initiates multifaceted inter-kingdom interactions that can fundamentally alter both parasite virulence and disease progression^6,7^. Central to this interaction is the parasite’s role as a microbial predator, which actively shapes community composition by preying on resident bacteria through phagocytosis^8^.

Previous investigations have established that *E. histolytica* utilizes specialized enzymatic mechanisms to circumvent the physical protection afforded by bacterial biofilms. Specifically, using *Bacillus subtilis* as a model for robust biofilm formation under gut-mimetic, nutrient-limited conditions, we demonstrated that amoebic cysteine proteinases (CPs) function as critical virulence factors that degrade the extracellular matrix via the proteolytic cleavage of the structural protein TasA ^9,10^. These findings further highlighted a significant divergence in the nature of inter-kingdom interactions depending on bacterial lifestyle: while CPs facilitate the mechanical breakdown of the sessile community^11^, the interactions of the parasite with planktonic populations appear to be governed by fundamentally distinct physiological and regulatory parameters.

The bacterial transition between motile, planktonic states and sessile, biofilm communities is a highly regulated behavioral switch driven by cyclic nucleotide second messengers^12^. While cyclic diguanylate (c-di-GMP) universally balances flagellar dynamics and extracellular matrix secretion, Firmicutes like *B. subtilis* integrate an additional, highly specialized regulatory layer through cyclic di-AMP (c-di-AMP) homeostasis ^13^. Cyclic di-AMP (c-di-AMP) regulates *Bacillus* growth, sporulation ^14^, DNA repair ^15^, osmoregulation ^16,17^, and biofilm formation ^18^. Cellular c-di-AMP levels are balanced by di-adenylate cyclase (DACs), which synthesize it from ATP, and specific degrading enzymes, directly modulating target protein activity^18^. These cyclic nucleotides allow the bacterium to rapidly translate microenvironmental stress cues into a definitive phenotypic switch, optimizing either spatial biofilm development or active defensive dispersal. However, while cyclic di-AMP (c-di-AMP) homeostasis plays a fundamental role in microbial adaptation to the environment, it has not been studied in the context of protist-bacteria interactions.

Adapted for the hypoxic environment of the human gastrointestinal tract, *E. histolytica* lacks mitochondria and a functional tricarboxylic acid (TCA) cycle, relying instead on a modified fermentative pathway for energy production^19–21^. A central feature of this metabolism is the bifunctional alcohol dehydrogenase 2 (EhADH2)^22^, which, alongside EhADH1 and EhADH3, facilitates the conversion of acetyl-CoA to ethanol via an acetaldehyde intermediate. Although these small molecules have traditionally been categorized as metabolic byproducts, their high diffusivity within the gut milieu suggests a broader role as inter-kingdom behavioral modulators. Given that these metabolites are known to generate autocrine chemokinetic responses in *E. histolytica*^23^, we hypothesized that *B. subtilis* might exploit these specific anaerobic signatures as a sensory cue to transition from a sessile biofilm to a motile, evasive phenotype, thereby circumventing predation before physical contact occurs.

To systematically decode this interkingdom signaling network, we used unbiased proteomic profiling of *B. subtilis* exposed to the secretome of the protozoan predator *Entamoeba histolytica*. We demonstrate that the amoebic secretome induces a coordinated, systems-level state transition in *B. subtilis*, reprogramming cells from a sessile biofilm state—anchored by extracellular matrix expression and low-to-intermediate c-di-AMP levels—to a motile, planktonic state. We identify acetaldehyde, a specific volatile byproduct of the amoebic alcohol dehydrogenase (EhADH) pathway, as the primary infochemical driving this cellular rewiring. Furthermore, while the capacity to integrate this metabolic cue into core regulatory circuits is conserved across natural isolates and diverse bacterial taxa, we show it is rapidly uncoupled during laboratory domestication. Together, these results provide a mechanistic framework for volatile-mediated interkingdom signaling, revealing how protozoan predators leverage metabolic flux to dictate bacterial decision-making and remodel gut community architecture.

## Materials and Methods

### Bacterial Strains and Media

All experiments were performed with *B. subtilis* NCIB 3610 ^24^. The list of strains is provided in Table S1. The strains were routinely manipulated in LB broth (Difco, Falcon BD, Corning, NY, USA) or LBGM medium (biofilm medium, in which manganese(II) sulfate was substituted with manganese chloride^25^) or MSgg medium (5 mM potassium phosphate, 100 mM MOPS pH 7, 2 mM MgCl_2_, 50 μM MnCl_2_, 700 μM CaCl_2_, 1 μM ZnCl_2_, 50 μM FeCl_3_, 2 μM thiamine, 0.5% glycerol, 0.5% glutamate, 50 μg ml^−1^ threonine, tryptophan and phenylalanine)^24^.

### Biofilm development assay

Cells from a single colony isolated on LB (Luria-Bertani) agar plates were grown to mid-logarithmic phase in 2 mL LB broth (4 h at 37°C with shaking). For plate biofilms, 2 µL of culture was spotted onto solid MSgg medium and incubated at 30 °C for ∼72 hours^26^. For floating biofilm assays, the inoculum ratio was 1:100. Pellicle phenotypes in cultures grown at 30°C were observed in 24-well plates (SPL Life Sciences, Korea)^27^. Images of colonies and pellicles were acquired with a Samsung ISOCELL GN1 sensor (50 MP, f/1.8 aperture, 1/1.31-inch sensor). Images were optimised for contrast and brightness, with adjustments kept consistent.

### Plate reader Assay

The *B. subtilis* strain used in this study is NCIB3610, which carries the *amyE*:P_sinI_-luciferase: cam gene. For all experiments, starter cultures were grown in LB Broth (BD Difco™ Dehydrated Culture Media: LB Broth, Thermo Fisher Scientific, Waltham, MA, USA). Cultures were diluted 1:1000 in defined MSgg medium and added to each well of a 96-well polystyrene Visiplate (Wallac), a white plate with a clear bottom. The cultures were then grown for 30 h at 30 °C in a Tecan Infinite 200 Pro plate reader (Tecan, Männedorf, Switzerland). We verified the experiment and the tool in two other plate readers, BioTek LogPhase 600 Microbiology reader and BioTek Synergy H1 Multimode Reader (Agilent Technologies, Santa Clara, CA, USA), and the optical density at 600 nm (OD_600_) and luminescence (with a sensitivity setting of 200) were measured every 20 min. The data in the figures represent averages from four replicate wells of a single representative experiment, where luminescence values were normalized to the highest recorded luminescence. All data were analysed using the lab-developed Mat-Tek software ^28,29^.

### Flow cytometry

*B. subtilis* 3610: *lacA::*P*_hag_*-gfp (cam) and *amyE*::P_tapA-tapA-sipW-tasA-_TasA-mCherry^30^ (cam) biofilms in liquid (static) were inoculated as described above and incubated for ∼14 hours. Biofilms were centrifuged, resuspended in PBS, and thoroughly vortexed. The samples were then mildly sonicated (SONICS VibraCell VCX750 sonicator), fixed in 4% paraformaldehyde (Thermo Fisher Scientific, USA), and stored at 4 °C until measurement. Flow cytometry was performed on a CytoFLEX S with manufacturer-default fluorescence detector setup for a violet laser (V450, 450/45 nm; V525, 525/40; V610, 610/20 nm and the 488 nm (blue) laser with the 525/40 BP filter (FITC/GFP channel) for P_hag_-gfp, 525/40 BP filter for P_tapA_-cfp.) and data acquisition using CytExpert software, including recording 100,000 events at a flow rate of ∼3500 events per second. We performed quality control using fluorescent (QC) beads according to the manufacturer’s protocol. For the negative control, we used a sample with no fluorescence. We performed extended cleaning cycles manually using Flow Clean reagent supplied by the manufacturer and double-distilled water at the beginning and end of each flow cytometry session, verifying a low event rate in filtered PBS ^26,31^.

### Confocal Microscopy

*B. subtilis* 3610, *lacA:*P*_hag_*-gfp (mls) and *amyE*: P_tapA-tapA-sipW-_TasA-mCherry (Cam) biofilms in liquid (static) were inoculated as described above and incubated for a time period of ∼14 hours^10,32^. Biofilms were centrifuged, resuspended in PBS, and then fixed in 4% paraformaldehyde. Images were captured with the STELLARIS 5 Confocal System (Leica Microsystems GmbH) using a White Light Laser (WLL) and a Diode UV laser, with HyDS spectral detectors, on a DMi8 inverted microscope. Hag-GFP (green fluorescence) was excited with WLL at 494 nm, and emission was recorded from 495–540 nm, indicating the presence of motile *hag*-expressing cells. Tap-mCherry (red fluorescence) was excited with WLL at 549 nm, and emission from 590–690 nm indicated the biofilm-forming cells expressing matrix TasA protein ^26^.

### Acetaldehyde Analysis

A high-performance liquid chromatography using a Thermo Scientific Vanquish HPLC system equipped with an autosampler. Data acquisition and processing were performed using Thermo Scientific Dionex Chromeleon Chromatography Data System (CDS) software (version 7.2, Thermo Fisher Scientific, Sunnyvale, CA / Waltham, MA, USA. Separation was performed on a Hypersil GOLD column (250 x 4.6 mm, 5 µ particle size) maintained at 25°C. The mobile phase consisted of acetonitrile and water (60:40, v/v) delivered isocratically at a flow rate of 1 ml min^-1^. Samples were filtered through a 0.22-micrometer membrane filter before injection, and 10 µl was injected for each run. Detection was performed at 289 nm.

We analyzed pure acetaldehyde under the same chromatographic conditions and used it as a retention-time reference. Amoeba spent medium samples were then analyzed using the same method. Peaks were detected and integrated into Chromeleon software. We assigned the acetaldehyde-associated peak in spent medium by matching its retention time to the pure acetaldehyde standard.

### Proteomics Analysis

#### Data-Independent Acquisition (DIA) Mass Spectrometry Data Processing

Raw Data-Independent Acquisition (DIA) mass spectrometry files (.mzML) were processed using DIA-NN (version 2.2.0) for peptide-spectrum matching, protein inference, and label-free quantification. To maximize identification rates and avoid database-induced bias, we generated an in silico spectral library directly from the *B. subtilis* subsp. subtilis 168 reference proteome (UniProt accession: UP000001570, 4,288 entries; downloaded September 10, 2025). We used the deep learning-based spectra and retention time predictor embedded in DIA-NN^33^. For multi-sample analyses, match-between-runs (MBR) was enabled alongside the ‘smart-profiling’ and ‘reanalyze’ options to maximize peptide identifications and minimize missing values across biological replicates. The search parameters were configured with a precursor-level false discovery rate (FDR) threshold of 1% (q-value 0.01) and a protein group-level FDR threshold of 1% (pg-level 1). Trypsin/P cleavage specificity was selected, allowing up to 2 missed cleavages. We restricted peptide length to 7–30 amino acids, precursor charge states to 1–4, and precursor m/z and fragment m/z ranges to 300–1800 and 200–1800, respectively. Protein group quantification was performed using the MaxLFQ algorithm based on fragment ion chromatogram extraction and integration^33,34^.

#### Data Preprocessing, Quality Control, and Imputation

Downstream data analysis was performed in Python (version 3.11) using the pandas (version 2.1), numpy (version 1.24), and scikit-learn (version 1.3) libraries. MaxLFQ protein group intensities were first log2-transformed. To ensure robust quantification and remove proteins with sporadic detection, we applied a stringent missing-value filter: we excluded protein groups with >= 50% missing values (NaN) across all samples, leaving a final matrix of 2,690 highly confident, quantified proteins.

We imputed missing values in the filtered matrix using a conservative, row-wise half-minimum approach. For each protein group, missing values were replaced with half of the minimum detected log2 intensity for that specific protein across all samples. This approach preserves the low-abundance signal of proteins that may be completely absent in certain conditions while avoiding the artificial variance reduction associated with global constant imputation^35^. Following imputation, systematic loading differences between mass spectrometry runs were corrected by median-centering each sample (subtracting the sample-specific median log2 from all values in that column)^34^.

#### Differential Expression Analysis

Pairwise differential expression (DE) analyses were performed comparing each of the three treatment conditions-Amoeba spent media, Acetaldehyde, and CBC-treated amoeba media-against the Control group. For each protein, a two-sided Welch’s t-test (unequal variance) was applied to the normalized, imputed log2 intensities. We selected Welch’s t-test over Student’s t-test to account for potential heteroscedasticity between the treatment and control groups. To control the family-wise error rate associated with multiple testing, we corrected raw p-values using the Benjamini-Hochberg (BH) false discovery rate (FDR) procedure. Proteins were classified as significantly differentially abundant if they satisfied a dual threshold of |log2 Fold Change| >= 2.0 (representing a >= 4-fold change in abundance) and a BH-adjusted p-value <= 0.05.

#### Dimensionality Reduction and Clustering

We performed Principal Component Analysis (PCA) on the normalized and imputed log2 intensity matrix using scikit-learn. The data were standardized (mean = 0, variance = 1) across proteins prior to singular value decomposition. PCA was performed on the full 12-sample dataset and a focused 9-sample dataset excluding the CBC condition to evaluate global sample clustering and identify the primary axes of experimental variation. Confidence ellipses (representing 1.5 standard deviations) were calculated and plotted for each sample group. We performed hierarchical clustering on the mean log2 intensities for each condition, derived from three biological replicates. We calculated distance matrices using either Euclidean or Pearson correlation distance (1 - r), then clustered them using Ward’s minimum-variance linkage. Finally, we generated heatmaps incorporating row annotations from SubtiWiki v5 alongside curated biological function groups^36^.

#### K-means Clustering

To group proteins into discrete, co-regulated expression patterns across all four conditions, K-means clustering was performed on the row-wise z-scored mean log2 intensities of the 2,690 proteins. Z-score standardization focused the clustering algorithm on expression-profile shape rather than absolute abundance. We determined the optimal number of clusters (k) using the elbow method, which identified k = 11 as the optimal inflection point for the within-cluster sum of squares (inertia). Mean expression profiles (mean +/− SD) were plotted for each of the 11 clusters to visualize the dominant proteomic trajectories^36^.

#### Functional and Regulon Enrichment Analyses

To identify the biological pathways and regulatory networks driving the observed proteomic shifts, three independent enrichment methodologies were applied to the lists of significantly upregulated and downregulated proteins: Sigma Factor Regulon Enrichment: *B. subtilis* Sigma factor regulon assignments were retrieved from the SubtiWiki v5 database^37^. For each of the 14 major sigma factors, over-representation within the significant up- and downregulated protein sets was evaluated using a one-sided Fisher’s exact test against the quantified proteome background. p-values were adjusted across all tested sigma factors within each condition using the BH FDR procedure.

GO and KEGG Pathway Enrichment: Gene Ontology (GO) terms were retrieved from the UniProt reference proteome GAF file, and KEGG pathway definitions for B. subtilis (organism code: bsu) were retrieved directly from the KEGG REST API. Locus tags (BSU numbers) were mapped to standard gene names to ensure exact matching. We assessed over-representation using a one-sided Fisher’s exact test, and corrected p-values using the BH FDR method. Terms with an adjusted p-value <= 0.05 and a minimum overlap of 2 proteins were considered significantly enriched^38^. PANTHER Over-representation Analysis: Significant protein lists were queried against the PANTHER database (taxon ID: 224308 for B. subtilis 168) using the PANTHER REST API. Over-representation was evaluated for GO Biological Process, GO Molecular Function, GO Cellular Component, and PANTHER Pathways using Fisher’s exact test with built-in FDR correction^39^.

#### Statistical Rigor of Proteomic Analysis

All statistical tests performed in this study were two-sided unless otherwise specified. We evaluated normality assumptions using Shapiro-Wilk tests where appropriate, and addressed heteroscedasticity by using Welch’s t-tests. Multiple testing corrections were applied globally across all features within each independent analysis using the Benjamini-Hochberg FDR procedure^35^.

### Crystal Violet assay

To evaluate the inhibitory effect of volatile acetaldehyde on *E. coli K-12* biofilm formation, Standard non-tissue-culture-treated polystyrene 24-well microtiter plates (Falcon, BD Biosciences) were used. The stationary-phase *E. coli* culture was diluted 1:100 in fresh TSB + 0.2% glucose to an initial optical density at 600 nm (OD600) of approximately 0.05. 1.0 mL of a freshly prepared 200 mM aqueous acetaldehyde solution (Sigma-Aldrich) was added to the first well, thereby establishing a volatile source. 1.0 mL aliquots of this cell suspension were added to the adjacent inoculum wells, creating a gradient from the volatile acetaldehyde source. Biofilm biomass was quantified using a modified microtiter dish crystal violet staining protocol. Following the incubation, the liquid culture containing planktonic cells was discarded by gentle inversion. The wells were washed three times with 1.5 mL of sterile phosphate-buffered saline (PBS, pH 7.4) to remove non-adherent and loosely attached bacterial cells, taking extreme care not to disrupt the adherent biofilm on the walls and bottom of the wells. The plates were then inverted and air-dried at room temperature for 30 min. To stain the adherent biofilm, 1.2 mL of a 0.1% (w/v) aqueous solution of crystal violet (Sigma-Aldrich) was added to each well to ensure complete coverage of the biofilm boundary, and the wells were incubated at room temperature for 15 min. The dye was discarded, and the plates were washed thoroughly by gentle immersion in a large reservoir of deionized water to eliminate excess unbound stain. The plates were inverted onto paper towels, vigorously blotted to remove residual water, and allowed to air-dry completely at room temperature for 2 h. To quantify the stained biofilm, the bound crystal violet was solubilized by adding 1.2 mL of 70% Ethanol. The absorbance was measured at 570 nm (OD570) using a microplate spectrophotometer ^40,41^.

## Results

### Amoebic secretome inhibits biofilm formation

To determine if *E. histolytica* produces metabolites that influence *B. subtilis* biofilm development, we assessed pellicle biofilm formation and *sinI* gene expression in the presence of *E. histolytica* lysate and secretions. Biofilm matrix expression is governed by the master regulator Spo0A, which activates transcription of the antirepressor *sinI*; SinI subsequently binds and inhibits the matrix repressor SinR ^42–44^. To evaluate whether E. histolytica secretions disrupt this pathway, we monitored pellicle biofilm formation and real-time *P_sinI_*-luciferase reporter activity in the presence of amoebic lysates or spent media.

Although there was pellicle biofilm formation in all the conditions, there was a delayed pellicle formation and an inhibition of the wrinkly architecture in the presence of 10% *E. histolytica* extracts (**Fig. 1A**) and 2% *E. histolytica* spent media (**Fig. 1B**). The P*_sinI_*-luciferase expression was reduced in the presence of *E. histolytica* lysate (**Fig. 1C**) and its spent media (**Fig. 1D**) at 10% and 1-2 %, respectively, indicating the secretions from *E. histolytica* affected the biofilm formation in *B. subtilis*. The peak activity, expression efficiency, fold change and total expression of P*_sinI_*-luciferase were reduced in 1-2% of Eh spent media and completely abolished in 5-10% concentrations of Eh spent media, whereas the expression time of P*_sinI_*-luciferase remained the same (**Fig. 1E**). These results indicate that *E. histolytica* releases soluble secreted effectors that potently suppress the transcription of genes required for biofilm matrix assembly.

**Figure 1.**
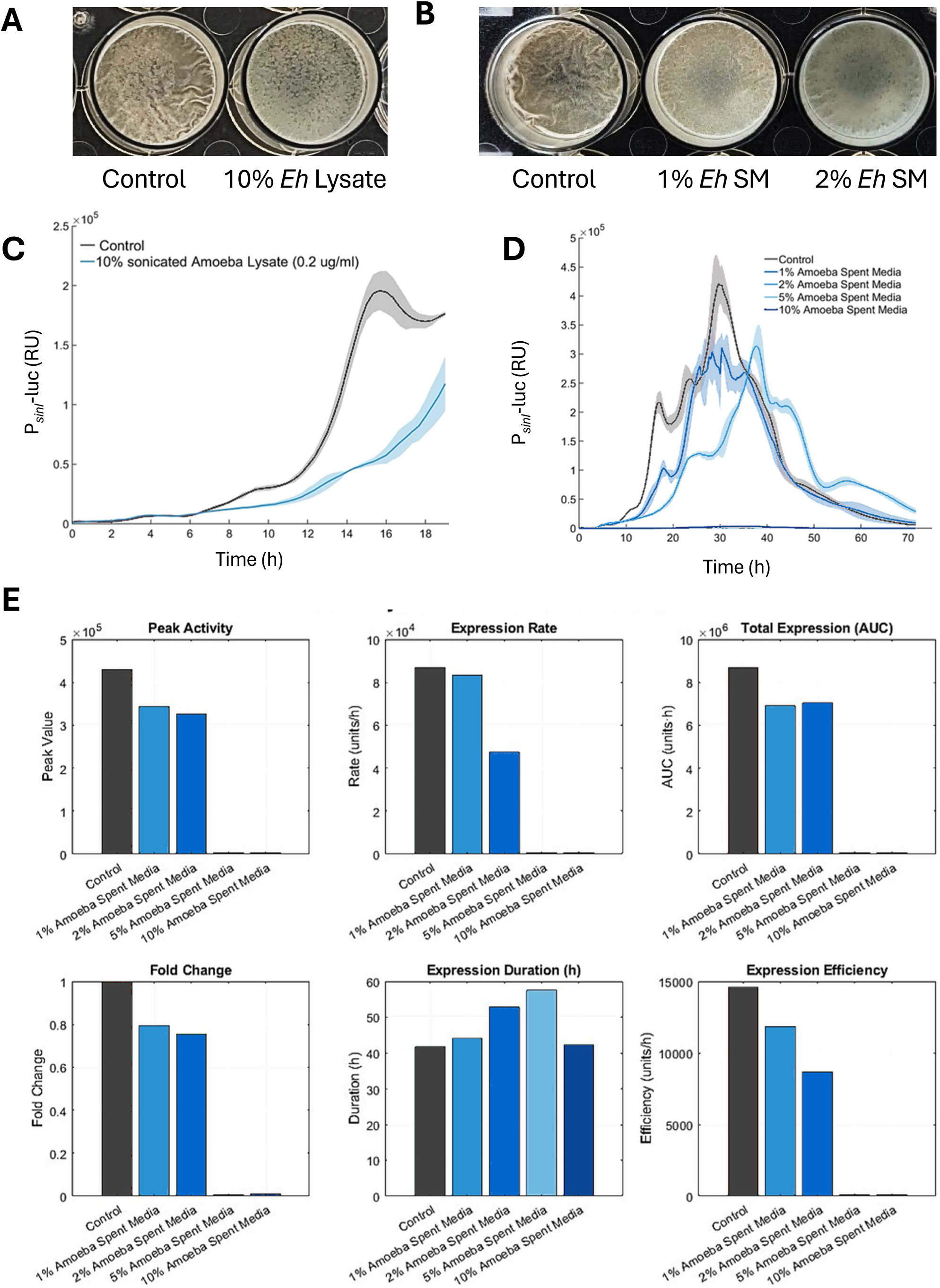
*E. histolytica* lysate and secreted factors suppress *B. subtilis* biofilm formation and *sinI* gene expression. *B. subtilis* pellicle biofilms formed in 24-well microtiter plates. **(A)** Biofilms grown in the absence (Control) or presence of 10% *E. histolytica* (*Eh*) lysate. **(B)** Biofilms grown in the absence (Control) or presence of 1% and 2% *E. histolytica* spent media. Analysis of *sinI* promoter expression using a P*_sinI_*-luciferase reporter strain. **(C)** Expression profile of P*_sinI_*-luciferase treated with 10% sonicated amoeba lysate compared to a control over time. Shaded regions indicate the standard deviation. **(D)** Dose-dependent effect of *E. histolytica* spent media (1%, 2%, 5%, and 10%) on P*_sinI_*-luciferase expression over time. **(E)** Quantitative analysis of kinetic parameters derived from the data in **(D)**. Bar graphs show Peak Activity, Expression Rate, Total Expression (Area Under the Curve, AUC), Fold Change, Expression Duration, and Expression Efficiency for the control and at different concentrations of *E. histolytica* spent media.

We next examined the effect of the *E. histolytica* secretions on *B. subtilis* behaviour by single-cell analysis. For flow cytometry analysis, we used two strains of wild-type *B. subtilis* harbouring *P_tapA_*-CFP (Cyan Fluorescent Protein) as a reporter of extracellular matrix (ECM) production and *P_hag_*-GFP (Green Fluorescent Protein) as a reporter of flagellar motility. For confocal microscopy analysis, we used a dual-labeled (TasA-mCherry and *P_hag_*-GFP) strain of *B. subtilis.* These strains were grown in MSgg media (Biofilm-inducing media) without shaking for ∼14 hours. In the presence of 2% *Eh* spent media, a lower percentage (3-fold decrease) of cells expressed the *P_tapA_*-CFP reporter than in the control (**Fig. 2A**). Concurrently, a higher percentage of cells expressed the *P_hag_*-GFP reporter than in the control (**Fig. 2B**). Using confocal microscopy, we found that cells were either expressing the ECM reporter (TasA-mCherry) or the motility reporter (*P_hag_*-GFP); neither was expressed at the same time (**Fig. 2C&D**). Compared with the control, *E. histolytica* lysates (**Fig. 2C**) or spent media (**Fig. 2D**) induced a significantly higher proportion of *B. subtili*s cells to activate motility, reaching 84% GFP-positive cells in spent media (**Fig. 2D**). The results (**Fig. 2A–D**) are consistent with the resolved biofilm and motility regulation pathways of *B. subtilis*, in which reducing *sinI* expression elevates levels of free SinR thereby repressing SlrR and preventing motility repression by the SinR•SlrR complex^45^, (**Fig. 2E**).

**Figure 2.**
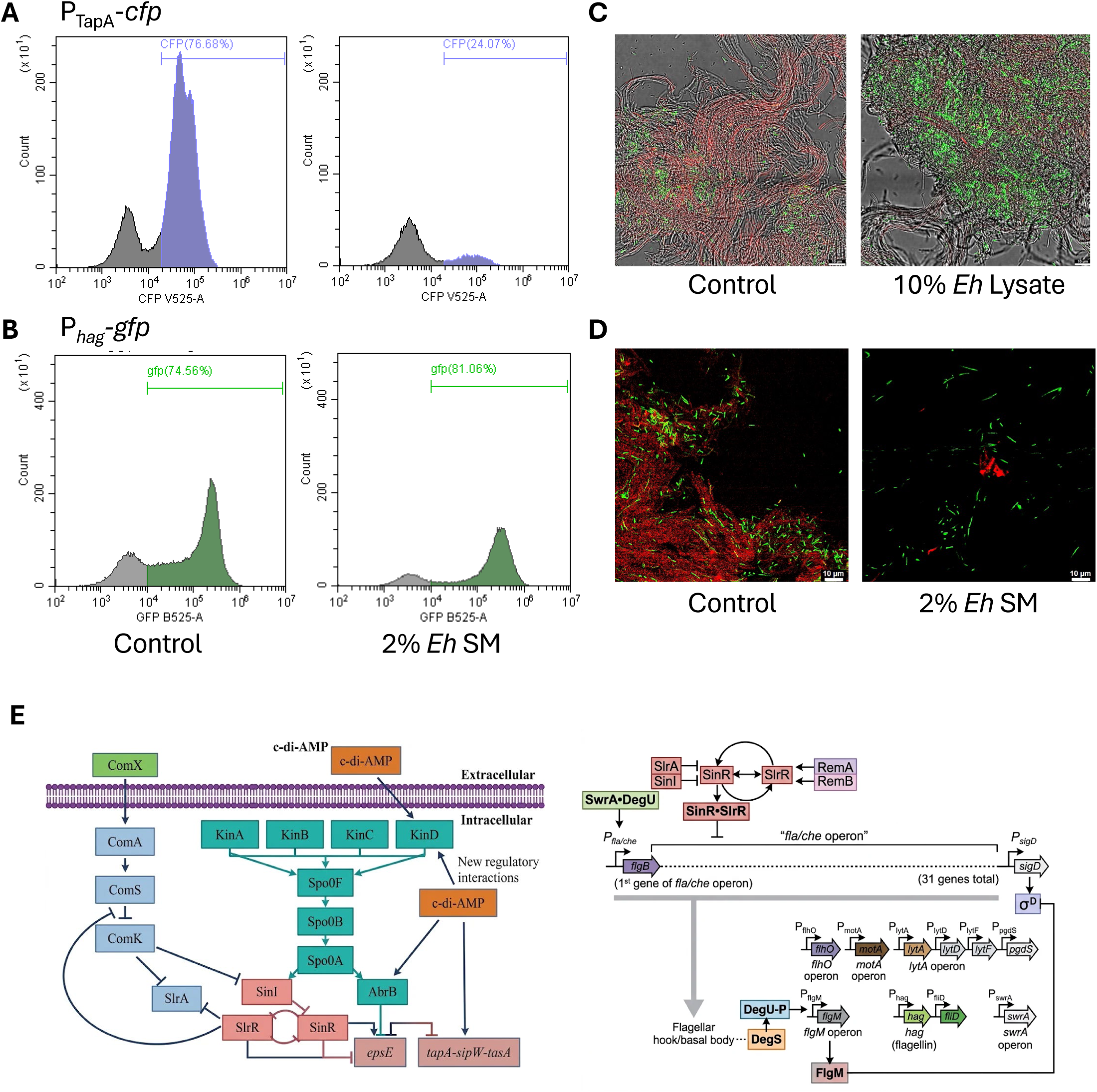
*E. histolytica* metabolites trigger a phenotypic switch towards motility in *B. subtilis*. **(A)** Flow cytometry analysis of matrix gene expression P*_tapA_*-CFP in *B. subtilis*. Histograms show the population distribution in control conditions (left) versus treatment with 2% *E. histolytica* spent media (right). The percentage of cells expressing the matrix gene drops from 76.68% to 24.07% upon treatment. **(B)** Flow cytometry analysis of flagellar gene expression P*_hag_*-GFP. Histograms compare control populations (left) to those treated with 2% *E. histolytica* spent media (right). The proportion of motile, flagellated cells increases from 74.56% to 81.06% in the presence of amoebic secretions. **(C-D)** Confocal laser scanning microscopy (CLSM) visualisation of the biofilm architecture: **(C)** Comparison of control biofilms (left) and those treated with 10% *E. histolytica* lysate (right). The control shows dense, matrix-rich structures (TasA-mCherry), whereas the lysate-treated sample shows disrupted architecture with an increase in individual motile cells (P*_hag_*-GFP). **(D)** Comparison of control biofilms (left) and those treated with 2% *E. histolytica* spent media (right). Treatment results in a loss of aggregated matrix-rich structures (TasA-mCherry) and an increase in individual motile cells (P*_hag_*-GFP). (**E**) **Biofilm and motility regulation pathways** (Left): Signaling network controlling biofilm matrix genes (*epsE* and *tapA-sipW-tasA*) via Spo0A, c-di-AMP, and the SinI/SinR/SlrR switch. (Right) Regulation of the *fla/che* operon and downstream *σ*^D^-dependent motility genes by the SinR•SlrR complex. Arrows denote activation; T-bars denote repression.

#### The amoebic EhADH metabolic pathway induces bacterial flagellar motility through the production of acetaldehyde

Because *E. histolytica* lacks mitochondria, it relies on anaerobic pathways for energy generation (**Fig. 3A**)^19–22^. Alcohol dehydrogenase 2 (EhADH2) is a bifunctional aldehyde/alcohol dehydrogenase that catalyzes the conversion of acetyl-CoA to ethanol through the intermediate formation of acetaldehyde. Although the parasite encodes three alcohol dehydrogenases (EhADH1, 2, and 3) with differential expression depending on environmental conditions, we focused on EhADH2 due to its central role in energy metabolism and its potential impact on parasite physiology and virulence^46^. We first explored the functional role of the ADH enzyme. To this end, we monitored *sinI* expression in *B. subtilis* using a *P_sinI_*-luciferase fusion reporter exposed to 5% *E. histolytica* lysate supplemented with ADH (300 U). The P*_sinI_*-luciferase expression was reduced in the presence of *E. histolytica* lysate (**Fig. 3B**), and the SinI reduction was enhanced by the addition of ADH. Treatment with ADH alone did not alter *sinI* expression. This result indicates that the signal is not ethanol, as it is degraded by ADH, but rather a possible product of ADH-mediated degradation.

**Figure 3.**
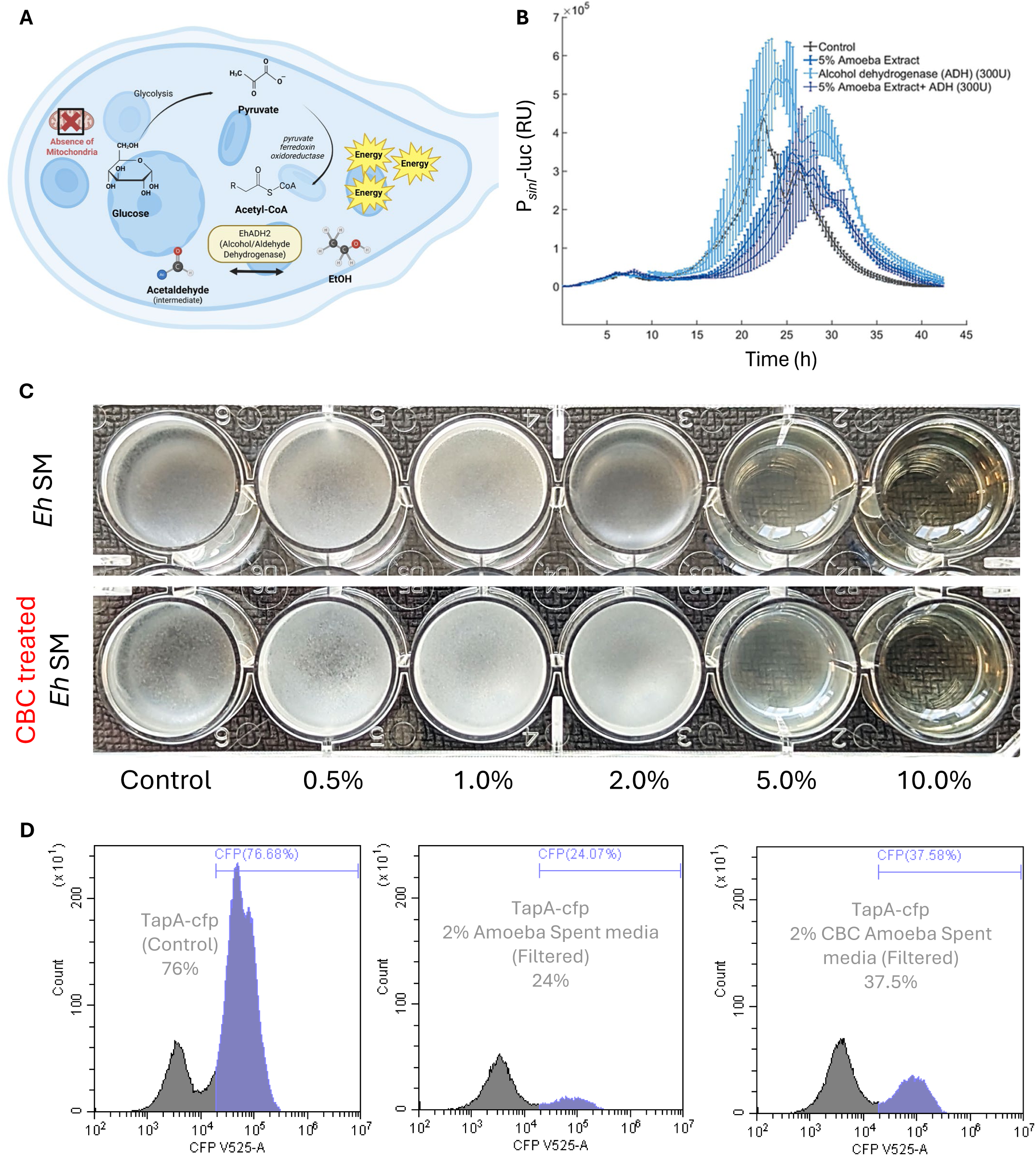
The inhibitory signal results from *E. histolytica* alcohol dehydrogenase (EhADH)-mediated fermentation. **(A)** Schematic illustration of *E. histolytica* energy metabolism. In the absence of mitochondria, the amoeba generates energy via fermentation, utilizing alcohol dehydrogenase (EhADH) to convert pyruvate to ethanol via acetaldehyde as an intermediate. **(B)** Kinetic analysis of *sinI* expression in *B. subtilis* P*_sinI_*-luciferase exposed to exogenous ADH with amoeba extract. Luminescence was monitored in cells treated with 5% *E. histolytica* lysate, purified ADH (300 U), or a combination of both, compared to a control. Data represent mean ± SD. **(C)** Biofilm phenotypes of *B. subtilis* treated with lysates from control amoebae versus amoebae treated with the ADH inhibitor Cyclobutyl Carbinol (CBC). Images show pellicle formation in 24-well plates with 10% lysate concentration. **(D)** Dose-dependent analysis of biofilm formation in the presence of Amoeba Spent Media (ASM). *B. subtilis* was cultured with increasing concentrations (0.5% to 10%) of spent media derived from either untreated or CBC-treated amoeba.

Taken together, these results suggest that metabolic products generated by EhADH activity contribute to inhibiting *B. subtilis* biofilm formation. To validate this hypothesis, we partially inhibited EhADH2 activity using Cyclobutyl Carbinols (CBC)^47^ and assessed the impact on *B. subtilis* behavior. We observed delayed pellicle formation in the presence of lysates and spent media derived from CBC-treated amoebae compared to untreated controls (**Fig. 3C**). These findings indicate that metabolites produced by the EhADH pathway modulate *B. subtilis* cells. We further investigated the effects of amoeba-secreted factors and the role of amoeba energy metabolism on *B. subtilis* using *P_tapA_-*CFP as a reporter and proteomic analysis. Flow cytometry results showed that in the control population, 76% of *B. subtilis* cells were positive for *P_tapA_-*CFP (**Fig. 3D**). Treatment with 2% *E. histolytica* spent media significantly reduced this proportion to 24%. However, when the amoeba was treated with CBC, an inhibitor of EhADH, the secreted factors in 2% CBC Amoeba spent media partially restored the *P_tapA_-CFP* positive population to 37.5%.

#### Acetaldehyde-mediated biofilm dispersal is conserved across undomesticated Bacilli but lost during domestication

After identifying ADH’s role in previous experiments, we tested whether the substrate (ethanol) or the ADH pathway intermediate, acetaldehyde, caused the observed inhibition. We first tested the effect of ethanol on *B. subtilis* biofilm formation and biofilm gene expression (**Fig. S1**). As shown in **Fig 4A**, the pellicle structure remained largely intact at lower ethanol concentrations up to 84 mM, and inhibition was observed only at high ethanol concentrations (up to 420 mM) as a solute and as a volatile (Liquid ethanol titration: Fig. 4A and Fig. S1A; volatile ethanol exposure: Fig. S1B), suggesting that ethanol may not be the primary inhibitory signal.

**Figure 4.**
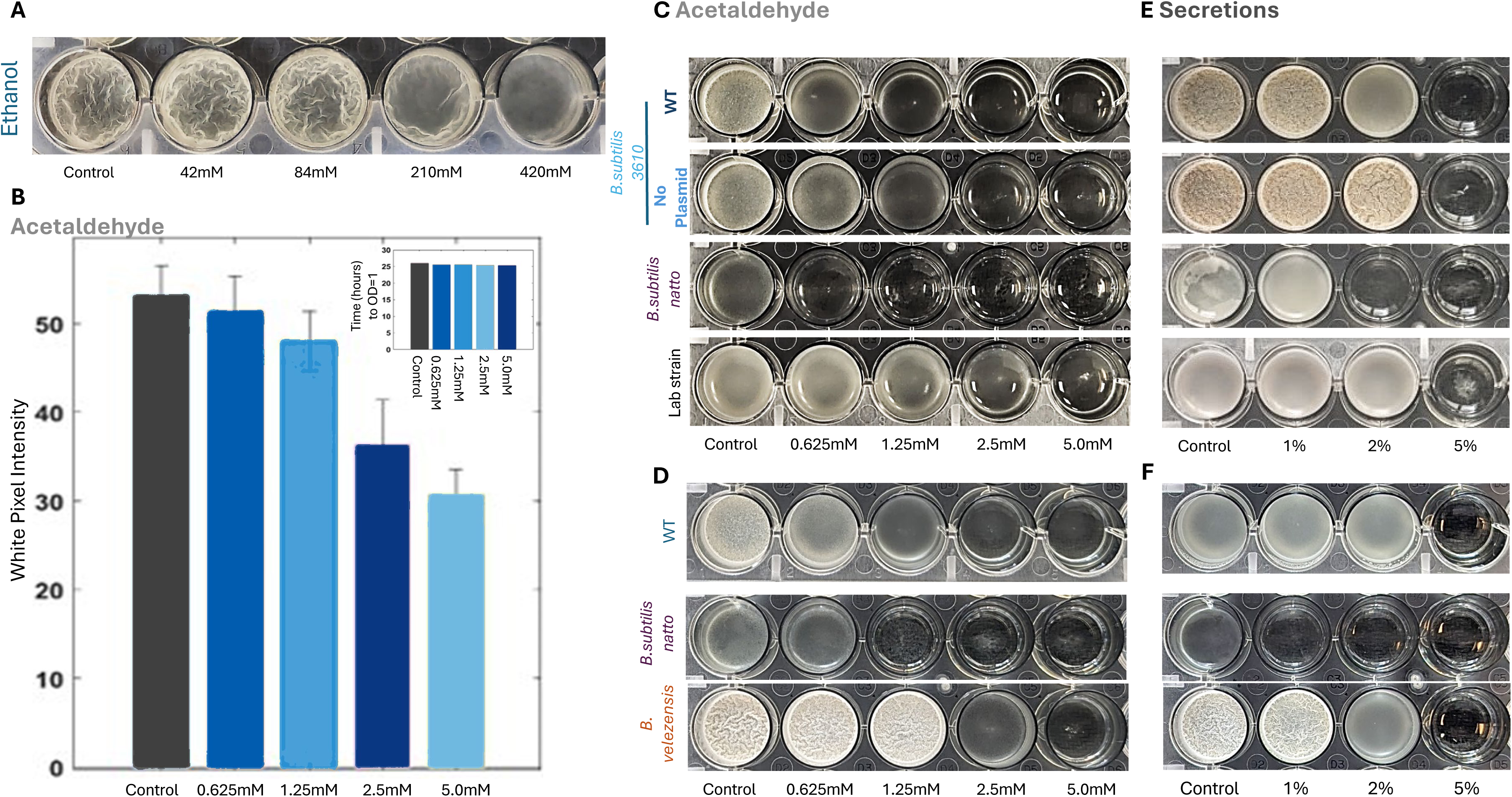
Acetaldehyde acts as the central signaling molecule for biofilm inhibition in *B. subtilis*. **(A)** Biofilm formation assay of *B. subtilis* in the presence of increasing concentrations of Ethanol (42 mM to 420 mM). Images show pellicle morphology in 24-well plates compared to a control. **(B)** Quantitative analysis of biofilm formation represented by white pixel intensity derived from the images in panel B. The data show a dose-dependent reduction in biofilm robustness with increasing Acetaldehyde concentrations, Insert: Growth kinetics of *B. subtilis* treated with Acetaldehyde. The bar graphs show the time required to reach an optical density (OD) of 1.0 at varying acetaldehyde concentrations (0.625 mM to 5 mM), indicating no significant growth inhibition. **(C)**. Pellicle Biofilm formation Assays in LBGM media at different concentrations of acetaldehyde of wild type (WT), B. subtilis WT strain without plasmid, B. subtilis natto, and laboratory strain 168. **(D)** Pellicle Biofilm formation Assays in MSgg media at different concentrations of amoeba spent media from wild type (WT), *B. subtilis* WT strain without plasmid pBS32, *B. subtilis* natto, and *B. subtilis* Laboratory strain**. (E)** Pellicle Biofilm formation Assays in LBGM media at different concentrations of acetaldehyde and (F) amoeba spent media from wild type (WT), B. megaterium, B. subtilis natto, and *B. velezensis*.

In contrast, we examined the effect of acetaldehyde, the metabolic product of ADH. A time-course analysis revealed an effective, dose-dependent inhibition of biofilm formation (**Fig. 4B**). While the control and those treated with low concentrations (0.625 mM) developed robust, wrinkled pellicles by 42 and 54 hours, cultures treated with 2.5 mM and 5.0 mM acetaldehyde exhibited a severe delay or failure in developing complex biofilm architecture. *B. subtilis* is influenced by the volatile compounds (VOCs) it produces, which help manage its population density. High concentrations of its own VOCs can suppress biofilm formation, leading to smaller, flatter colonies without their typical 3D structure by reducing the expression of extracellular matrix genes^48^. Additionally, *B. subtilis* can detect external microbial volatiles^49^. Therefore, we tested whether acetaldehyde could serve as a volatile. Using a volatile exposure assay that physically separated the volatile source from the bacterial culture, we found that acetaldehyde significantly inhibited *B. subtilis* pellicle formation without direct contact (**Fig. S2**). These results show that acetaldehyde retains biological activity in the gas phase and suggest it may function as a diffusible, long-range signaling molecule that modulates bacterial biofilm development.

To confirm that the lack of biofilm resulted from specific signaling interference rather than cellular toxicity, we analyzed the growth kinetics of *B. subtilis* in the presence of acetaldehyde. We used a plate reader to analyze growth curves and determine the time required for cultures to reach ODs of 1.0 (Mid-Log) (**Fig. 4B**). The results demonstrated that the time to reach these growth milestones was comparable between the control and all tested acetaldehyde concentrations, confirming that the metabolite inhibits biofilm development without impeding planktonic growth. Chromatographic analysis further confirmed acetaldehyde in the protist growth medium (**Fig. S3**).

To determine whether this acetaldehyde-mediated signaling is conserved across the *Bacillus* genus, we next compared the responses of wild, environmental isolates with a domesticated laboratory strain. This comparison revealed a clear phenotypic divergence. Undomesticated, robust biofilm-forming isolates—including *B. subtilis* 3610^50^, *B. subtilis* var. *natto* (known for its tenacious, viscous extracellular matrix)^50^, and the plant-associated colonizer *B. velezensis*^51^—were highly sensitive to both purified acetaldehyde (**Fig. 4C, D**) and *E. histolytica* cues from spent media (**Fig. 4E, F**). Across these wild strains, exposure drove pronounced, concentration-dependent reductions in air–liquid interface pellicle formation and promoted the transition to motility. In stark contrast, the domesticated reference strain *B. subtilis* 168^52^ was completely unresponsive to both acetaldehyde (**Fig. 4C**) and *E. histolytica* spent media (**Fig. 4E**), exhibiting neither motility induction nor signal-mediated disruption of pellicle assembly. Interestingly, this response is partially regulated by the large 84-kb plasmid pBS32 in *B. subtilis* NCIB 3610. pBS32 encodes the phosphatase RapP, a known inhibitor of biofilm formation that dephosphorylates Spo0F∼P to dampen the Spo0A phosphorelay. Consistent with this repressive role, curing pBS32 ^53,54^ rendered NCIB 3610 more resistant to amoeba-derived signals (**Fig. 4C** and **D**).

#### System-Wide Proteomic Reprogramming of B. subtilis by Amoeba Cues and Acetaldehyde

To determine the genetic basis of acetaldehyde-induced inhibition, we evaluated its effects on *B. subtilis* strains lacking key biofilm regulators. Interestingly, the wild-type (WT) strain showed a dose-dependent sensitivity in pellicle formation to acetaldehyde, whereas mutants for the master regulators *abrB* and *sinR* maintained robust pellicle formation even at high concentrations (up to 5 mM) (**Fig. 5A**). The matrix-deficient mutant Δ*epsH* showed no significant change in pellicle formation (**Fig. 5A**). We further visualized the transition to motility gene expression in the presence of acetaldehyde using confocal microscopy in a dual-labelled (TasA-mCherry and P_hag-_GFP) strain of *B. subtilis*. The cells were grown in MSgg medium (biofilm-inducing medium) without shaking for ∼14 hours. Under normal conditions, *B. subtilis* cultures were characterized by dense clusters of matrix-producing red cells (TasA-mCherry). Upon treatment with acetaldehyde, these clusters were largely abolished, replaced by a dispersed population of highly motile cells expressing GFP (*P_hag_*-GFP) (**Fig. 5B**).

**Figure 5.**
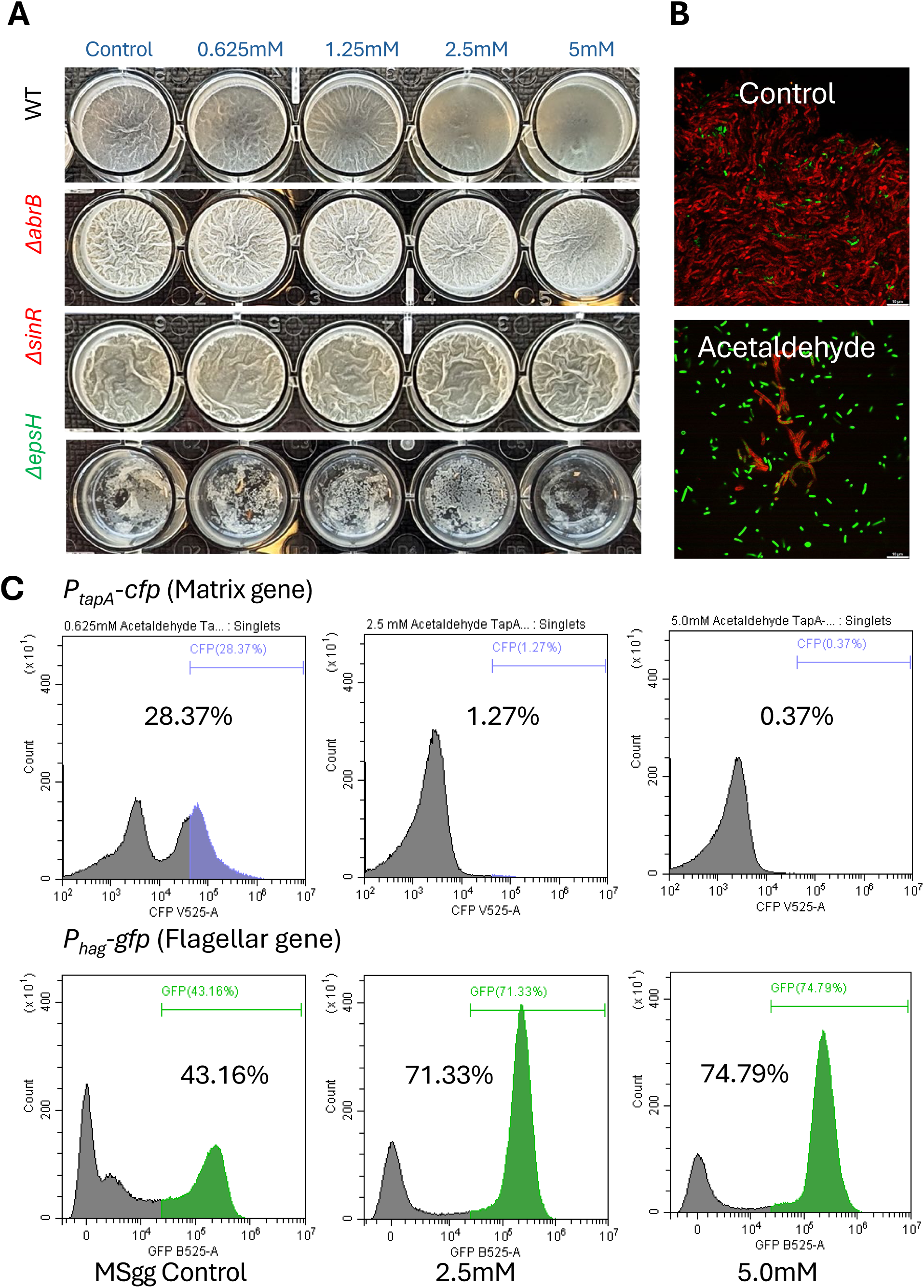
Acetaldehyde induces a transcriptional switch towards active motility in *B. subtilis*. **(A)** Pellicle biofilm formation assays of wild type (WT), biofilm regulator mutants Δ*abrB*, Δ*sinR*, and matrix repressor mutant Δ*epsH* across a gradient of acetaldehyde concentrations (0.625 mM to 5 mM). **(B)** Confocal microscopy images showing the distribution of matrix-producing cells (TasA-mCherry) and motile cells (P*_hag_*-GFP) in control versus acetaldehyde-treated conditions. **(C)** Flow cytometry histograms of P*_tapA_*-CFP (matrix gene) and P*_hag_*-GFP (flagellar gene) expression in MSgg control medium and under treatment with 2.5 mM and 5.0 mM acetaldehyde. Percentages indicate the fraction of the population positive for the respective fluorescent reporters.

This phenotypic shift was confirmed at the transcriptional level through flow cytometry using *P_tapA_*-CFP and *P_hag_*-GFP *B. subtilis* reporter strains. Treatment with 5.0 mM acetaldehyde decreased the matrix-producing subpopulation, with *P_tapA_*-CFP expression reducing from 29.33% in the control to 0.38%. Simultaneously, the proportion of cells expressing the flagellar gene P*_hag_*-GFP increased significantly, from 43.16% to 74.79%. This further confirms that acetaldehyde from Eh acts as a specific signaling molecule that transcriptionally reprograms *B. subtilis* to Favor a motile lifestyle over biofilm development (**Fig. 5C** and **2E**).

The specificity of acetaldehyde-related biofilm signals was significant, as ethanol did not cause a notable reduction in sinI expression up to 2.5% v/v (210 mM) (**see Fig. S4**). However, ethanol did play a role in biofilm regulation at high concentrations (5.0% v/v; 420 mM) (**see Fig. S1, an d S4**). The effect was also observed without direct contact using volatile ethanol exposure (**Fig. S1**). This effect was not observed in the *sinR* and *abrB* mutants, which lack the repressors of the extracellular matrix (**see Fig. S5**). These results were consistent with ethanol shifting cells toward motility (**Fig. S6)**, supporting a broader role for the amoeba’s fermentation signaling in inducing microbial motility. Although up to 5 mM acetaldehyde delayed pellicle formation and prevented wrinkly architecture, PsinI-luciferase expression remained stable, suggesting that motility regulation dominated over sinI expression. Ethanol (at high concentrations) and acetaldehyde influence the same bacterial sensory pathway, as their combined effects on biofilm formation and *sinI* promoter were additive (**Fig. S7 and S8**), and both were dependent on SinR and AbrB. This finding aligns with our genetic findings that intact *sinR* and *abrB* regulatory circuits are strictly required for signal perception, as domesticated strains like 168 harbor mutations ^52^ that attenuate these master regulators.

To characterize how *B. subtilis* responds to amoeba-derived factors, we performed comparative proteomic profiling of untreated cells versus those exposed to either amoeba spent media or acetaldehyde (**Fig. 6A** and **Fig. S9**). Amoeba media induced extensive proteomic remodeling: proteins involved in flagellar assembly, chemotaxis, two-component signaling, and teichoic acid biosynthesis were significantly upregulated (**Fig. 6A, S9 and S10 for gene ontology enrichment**), while arginine biosynthesis and other discrete metabolic pathways were downregulated. This indicates a coordinated physiological reorganization rather than nonspecific metabolic inhibition. Because we previously identified acetaldehyde as a major inhibitory metabolite in this interaction, we investigated its specific contribution. Strikingly, acetaldehyde treatment recapitulated core signatures of the spent-media response, including the induction of motility apparatuses, the suppression of biofilm-associated, Spo0A-regulated pathways (F**ig. 6B****, S9 and S10**), and broader cellular reorganization of small-molecule metabolism. Together, these findings establish acetaldehyde as a primary chemical driver that suppresses biofilm programs and stabilizes motility. Biofilm regulation is also manifested in the response of secondary messengers to exposure to the amoeba-spent medium. This exposure selectively rewires cyclic di-AMP (c-di-AMP) homeostasis, leaving the canonical cyclic di-GMP (c-di-GMP) network unperturbed (**Fig. 6C**). Specifically, biofilm formation seemed to correlate with a strong upregulation of the phosphodiesterase GdpP and concurrent downregulation of the diadenylate cyclase DisA. Because basal c-di-AMP pools actively restrain premature biofilm dispersal, this depletion relieves the antagonism and depresses flagellar genes, initiating evasion. These signaling dynamics collectively align with the heightened sensitivity observed in the *cdaA* mutant, which is defective in c-di-AMP synthesis (**Fig. 6D**).

**Figure 6.**
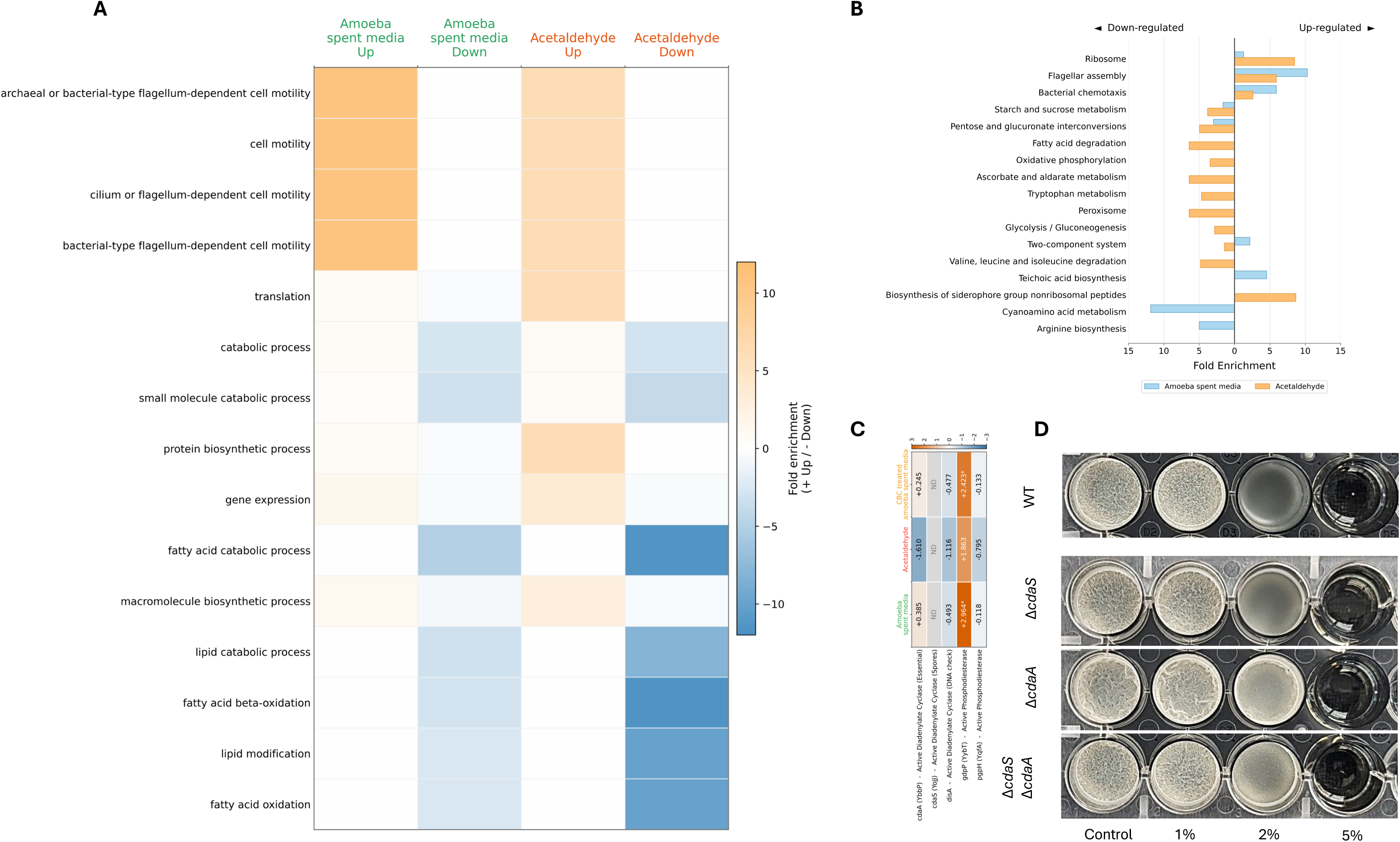
Global proteomic remodeling of *B. subtilis* following exposure to amoeba spent media and acetaldehyde. **(A)** Principal component analysis (PCA) of label-free quantification (LFQ) proteomics profiles from Control, Amoeba spent media, and Acetaldehyde-treated *B. subtilis*. Each point represents an individual replicate. PC1 explains 42.5% of the total variance and separates Acetaldehyde-treated samples from Control, and Amoeba spent media samples, whereas PC2 explains 39.8% of the variance and separates Amoeba spent media from Control. **(B)** Bidirectional KEGG pathway bar plot showing the number of overlapping proteins assigned to enriched KEGG pathways. Bars extending to the right indicate enrichment among upregulated proteins, whereas bars extending to the left indicate enrichment among downregulated proteins. Blue bars represent the Amoeba spent media, and green bars represent the Acetaldehyde. **(C)** Gene Ontology biological process enrichment matrix for the same comparisons. Enrichment patterns highlight treatment-specific proteomic responses, including motility-associated processes in Amoeba spent media and metabolic or lipid-associated processes in Acetaldehyde-treated cells. Color scale represents fold enrichment, with red denoting enrichment in upregulated proteins and blue denoting enrichment in downregulated proteins. **(D)** Proteomic expression analysis of cyclic di-AMP (c-di-AMP) synthesis (*cdaA*, *cdaS*, *disA*) and degradation (*gdpP*, *pgpH*) proteins in *B. subtilis* exposed to amoeba spent media, acetaldehyde, or CBC-treated amoeba spent media. Heatmap values represent log_2_ fold changes relative to untreated controls. Asterisks denote statistically significant changes (*p < 0.05). ND = not detected.* **(E)** Effect of increasing concentrations of exogenous amoeba secretions on the pellicle biofilm formation of *B. subtilis* wild-type (WT) and c-di-AMP signaling mutants (Δ*cdaS*, Δ*cdaA*, Δ*cdaS*Δ*cdaA*) grown in MSgg medium.

While acetaldehyde alone triggers this localized, motility-driven escape, the full amoeba secretome induces a systemic, multi-layered defensive program that further enhances motility (**Fig. 7A and S10**). Profiling revealed a ∼2.5-fold enrichment in general stress-response proteins for environmental cross-protection and a ∼3.2-fold enrichment of the SOS/DNA repair pathway (including key proteins RecO and UvrA), suggesting the secretome induces significant DNA damage or oxidative stress. Alongside activating these defenses, *B. subtilis* aggressively represses nonessential developmental and metabolic pathways to maximize short-term survival. Specifically, urease activity and cell wall macromolecule catabolism are each suppressed by more than 20-fold. Halting peptidoglycan turnover likely reinforces the cell envelope against amoebic enzymatic degradation, while repressing urease shifts the cell away from routine nitrogen assimilation to conserve vital resources during acute stress.

**Figure 7.**
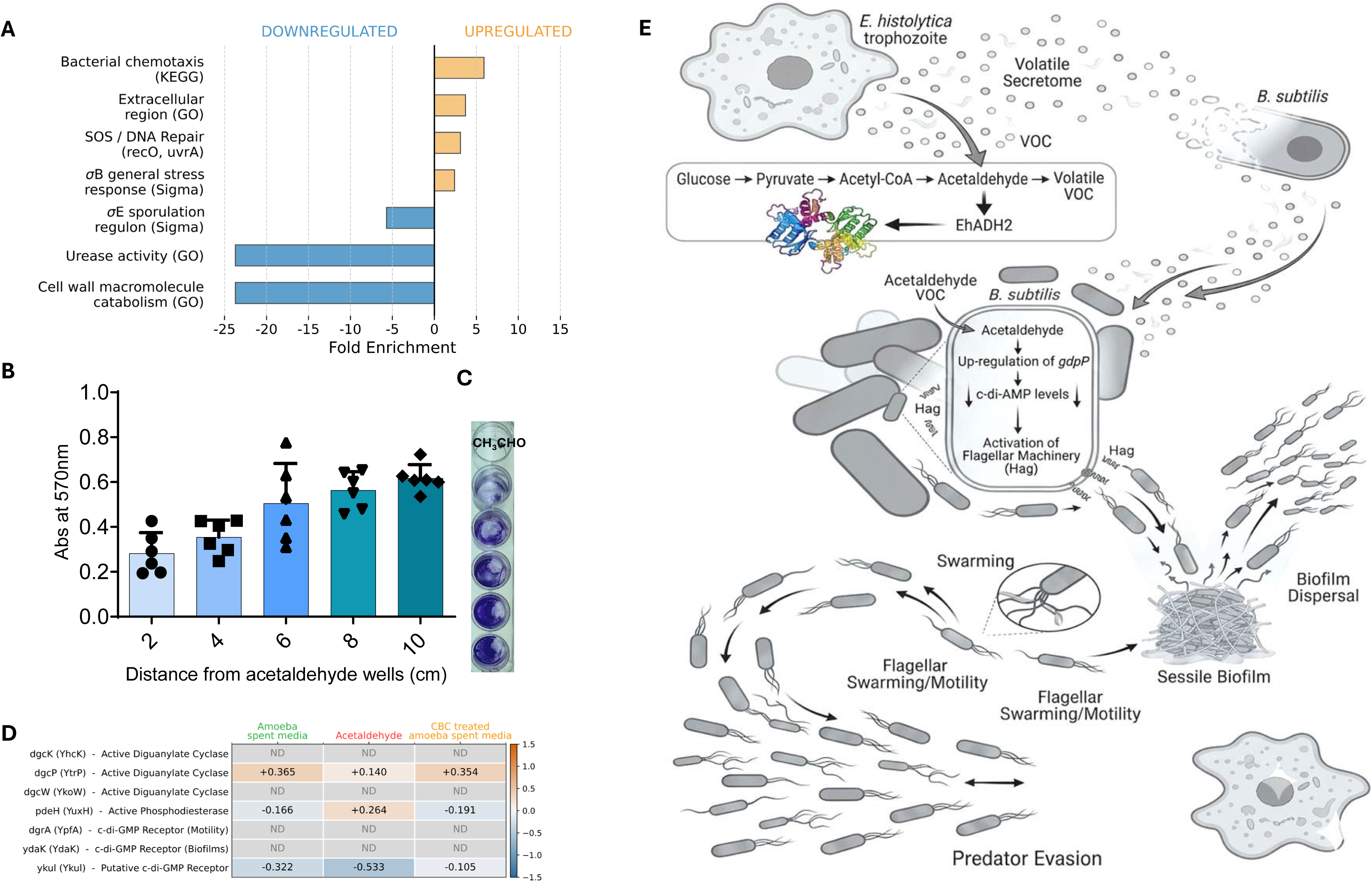
Cross-kingdom signaling responses across amoeba-derived and acetaldehyde-associated conditions. **(A)** Functional enrichment analysis of differentially expressed features, showing pathways or gene sets enriched among downregulated and upregulated responses. Negative fold-enrichment values indicate enrichment among downregulated features, whereas positive values indicate enrichment among upregulated features. Downregulated categories included urease activity, cell wall macromolecule catabolism, and the σE sporulation regulon; upregulated categories included bacterial chemotaxis, extracellular-region genes, SOS/DNA repair genes, and the σB general stress-response regulon. (**B, C**) Biofilm formation in *E. coli* in response to acetaldehyde as a volatile. Crystal violet is shown as the mean of 2 biological replicates (n = 3; left), with the corresponding plate (right) and an image of a representative experiment. **(D)** Heatmap of log_2_ fold changes for genes associated with c-di-GMP synthesis and degradation across the same conditions. Color intensity denotes log_2_ fold change relative to the corresponding control, with orange indicating increased expression and blue indicating decreased expression. Numerical values are shown within each cell; ND indicates not detected. (**E**) A model summarizing the amoeba-*B. subtilis* interactions

To determine whether this defensive sensing extends beyond Gram-positive *Bacilli,* we examined *Escherichia coli*, a Gram-negative prey of *E. histolytica* whose biofilms are similarly vulnerable to degradation by amoebic cysteine proteinases ^10^. Acetaldehyde alone was sufficient to inhibit *E. coli* biofilm formation, even when delivered strictly as a volatile organic compound (VOC) (**Figs. 7B** and **7C**). This demonstrates that evolutionarily distant bacterial phyla co-opt shared metabolic cues to detect and evade protozoan predation.

Cyclic dinucleotides play a conserved role in governing the switch between sessile biofilm formation and planktonic motility^12^ and we accordingly observed that c-di-GMP turnover is also significantly altered by acetaldehyde (**Fig. 7D**). We therefore hypothesize that these second messengers serve as a broadly conserved signaling hub orchestrating bacterial anti-predator responses across diverse taxa.

## Discussion

### Acetaldehyde as a Regulatory Infochemical in the Parasite–Microbiota Interface

*Entamoeba histolytica* remains a leading cause of global morbidity^55^, particularly in regions where the gut microbiome’s composition significantly dictates the transition from commensalism to invasive amoebiasis^2,56^. Our findings identify the ethanol degradation pathway-and specifically its volatile intermediate, acetaldehyde-as a primary metabolic mediator in this inter-kingdom dialogue (**Figs 2-3**). While acetaldehyde is a ubiquitous fermentation byproduct in mucosal environments, its role has traditionally been viewed through the lens of metabolic flux rather than regulatory signaling^57^. Acetaldehyde inhibits the yeast-to-hypha conversion and biofilm formation in *Candida albicans*^58^. By extending this concept to predator–prey dynamics, we show that *E. histolytica* uses this diffusible cue to fundamentally reprogram bacterial developmental decisions, stabilizing the prey microbiota in a dispersed, planktonic lifestyle.

### The Duality of Acetaldehyde: Signal and Stressor

The formation of biofilms in the presence of *E. histolytica* is likely to reduce the fitness of *B. subtilis* as the protist secretes cysteine proteinases (CPs) to proteolytically cleave structural matrix proteins, physically compromising the biofilm’s integrity^10,31^. Therefore, for planktonic counterparts, the metabolic byproduct acetaldehyde acts as a signal that upregulates motility and downregulates the matrix biosynthesis genes in Spo0A-regulated pathways (**Figs 4-6**) to increase the probability of successful migration to a site with low predator abundance. Beyond its role as a behavioral cue, acetaldehyde is a highly reactive electrophilic metabolite^59^. Its ability to form covalent adducts with proteins and nucleic acids suggests a deeper layer of physiological reprogramming. In eukaryotic systems, such modifications can alter chromatin architecture and transcriptional output; analogously, acetaldehyde may influence *E. histolytica* autocrine signaling by modifying its own regulatory proteins.

For bacterial prey, acetaldehyde likely serves as a dual-mode stimulus. At lower concentrations, it serves as a potent signaling molecule for the biofilm-to-motility switch. At higher concentrations, it may impose significant electrophilic stress, triggering conserved detoxification systems like AldA (aldehyde dehydrogenase)^60,61^. This overlap between stress response and developmental regulation highlights how *B. subtilis* has evolved to link its redox homeostasis to predatory avoidance, ensuring survival in the chemically volatile gut environment.

Ultimately, this phenotypic divergence reflects the distinct evolutionary pressures bacteria face in nature versus the laboratory. *E. histolytica* releases acetaldehyde as a specific signaling molecule that transcriptionally reprograms wild-type *Bacillus* strains—such as 3610, *B. subtilis var. natto,* and *B. velezensis (*up to 77% an Average nucleotide identity versus *Bacillus subtilis*^62^)—to abandon biofilm architecture via SinR-and AbrB-dependent pathways and favor motility, whereas the domesticated laboratory strain 168^52^ remains entirely resistant. Wild and environmental strains must maintain intact SinR- and AbrB-dependent regulatory pathways to escape protozoan grazing or metabolic stress by switching from biofilms to motility, whereas domesticated laboratory strains like 168 have lost these protective responses. Free from natural predators and environmental competition in culture, laboratory strains benefit from locking into a stable multicellular state without wasting energy on costly lifestyle transitions triggered by metabolic byproducts like acetaldehyde.

Crucially, our results demonstrate that acetaldehyde maintains its potency as a VOC (**Fig. S2**). In the complex micro-ecology of the human gastrointestinal tract—characterized by varying oxygen gradients and physical barriers-the ability to sense volatile signals provides a distinct evolutionary advantage^49^. Unlike contact-dependent signals, acetaldehyde diffuses rapidly, providing a “chemical preview” of an approaching predatory front. This distal sensing allows the bacterial population to preemptively transition to a motile phenotype, thereby navigating away from the “grazing” front of the amoeba’s directed-avoidance behavior (**Fig. 7D**).

### Multi-Modal Defensive Networks Shape the Bacterial Response to the Complex Secretome

In complex microbial ecosystems, prey organisms rarely rely on a single cue to navigate dynamic trophic interactions; instead, they must integrate multi-layered chemical information that distinguishes distant threats from imminent physical contact. While the isolated predator-derived volatile acetaldehyde functions as a precise, long-range proxy that triggers an anticipatory “flight” response, the complete amoeba secretome elicits a far more comprehensive, multi-layered physiological overhaul. As revealed by our functional enrichment profiles (**Figs. 6** and **7**), exposure to the full spectrum of amoeba secretions forces *B. subtilis* to pivot from simple behavioral avoidance to a robust and orchestrated defense aimed at preserving cell envelope integrity and genomic stability. Specifically, the pronounced enrichment of the general stress response alongside the SOS/DNA repair machinery— including key recombinational and nucleotide excision repair proteins such as RecO and UvrA—demonstrates that the broader predatory cocktail imposes severe oxidative and genotoxic stress^63^. Thus, *B. subtilis* employs a tiered sensory strategy: utilizing volatile metabolites as early-warning signals to disperse before contact, while simultaneously mobilizing high-cost cellular fortification and DNA repair programs to survive direct chemical assault during grazing.

In parallel with these protective investments, the *B. subtilis* regulatory network executes a profound, systems-level reallocation of cellular resources, shutting down nonessential developmental and metabolic pathways to prioritize short-term survival. This global economic shift is highlighted by an extreme, greater-than-20-fold downregulation of both urease activity and cell wall macromolecule catabolism. By severely restricting its own peptidoglycan turnover, *B. subtilis* structurally reinforces its cell envelope^64^, making the macromolecular barrier less susceptible to enzymatic exophagy or mechanical breach by amoebic cysteine proteinases (CPs). Simultaneously, the abrupt dampening of urease activity—tightly linked to biofilm biomineralization^65,66^—signals a strategic shift away from routine nitrogen assimilation. This coupled repression effectively conserves vital metabolic currencies, redirecting energy away from sessile developmental programs to fuel the acute transcriptomic demands of the SOS response and flagellar assembly.

### A Conserved Cyclic Nucleotide Hub for Predator Evasion

Our findings show that the predatory escape response of *B. subtilis* is orchestrated by a highly selective rewiring of the cyclic di-AMP (c-di-AMP) homeostasis network, which functions as a central regulatory hub. Rather than exhibiting global fluctuations in the canonical cyclic di-GMP (c-di-GMP) signaling pathway, exposure to amoeba-spent medium drives a stark regulatory asymmetry within the c-di-AMP circuit. The bacteria significantly upregulate the primary phosphodiesterase GdpP (+2.964) while concurrently downregulating the vegetative diadenylate cyclase DisA (−0.493). This reciprocal target modulation shifts the cellular equilibrium toward rapid intracellular c-di-AMP degradation. By funneling diverse environmental and metabolic predator cues into this single-nucleotide bottleneck, the system rapidly relieves transcriptomic repression of downstream flagellar machinery and licenses active motility, demonstrating a highly optimized decision-making architecture. Strikingly, this secondary messenger switch is conserved enough to span broad phylogenetic boundaries, including Gram-negative enterobacteria. As shown on **Fig. 7**, exogenous volatile acetaldehyde exposure is similarly sufficient to disrupt biofilm architecture and inhibit matrix development in *Escherichia coli*. Because cyclic nucleotides serve as universal, ancestral toggles governing the sessile-to-motile transition across both Gram-positive and Gram-negative cell walls, they represent an ideal, evolutionarily optimized hub for processing volatile predatory info-chemicals.

### Concluding Remarks

This study establishes the parasite-derived volatile acetaldehyde as a critical, previously unrecognized info-chemical operating at the *Entamoeba histolytica*–microbiota interface. By directly linking the unique metabolic fluxes of anaerobic protozoan fermentation to bacterial regulatory networks, these findings reinforce an emerging paradigm in which low-molecular-weight microbial byproducts function not merely as metabolic waste, but as sophisticated, highly diffusible agents of inter-kingdom communication.

Furthermore, clarifying how these chemical cues interact with conserved second-messenger cascades and multimodal stress responses offers key insights into the evolutionary arms race that governs predator-prey dynamics within the gut. Ultimately, deciphering these metabolic parameters provides a novel conceptual framework for targeted therapeutic interventions to modulate the chemical microenvironment of the intestinal niche, thereby safeguarding the architecture of the commensal microbiome against parasitic invasion and pathogenesis.

