## Supporting Figures and Legends for "Protozoan Anaerobic Metabolism Emits Acetaldehyde as a Volatile Infochemical Driving Bacterial Biofilm Dispersal"

### Deciphering the Signaling Network Underlying Bacterial Escape from Protozoan Predators

#### Supporting Information

#### Supporting Figures (S1-S10)

#### Supporting Table S1

#### Supporting References

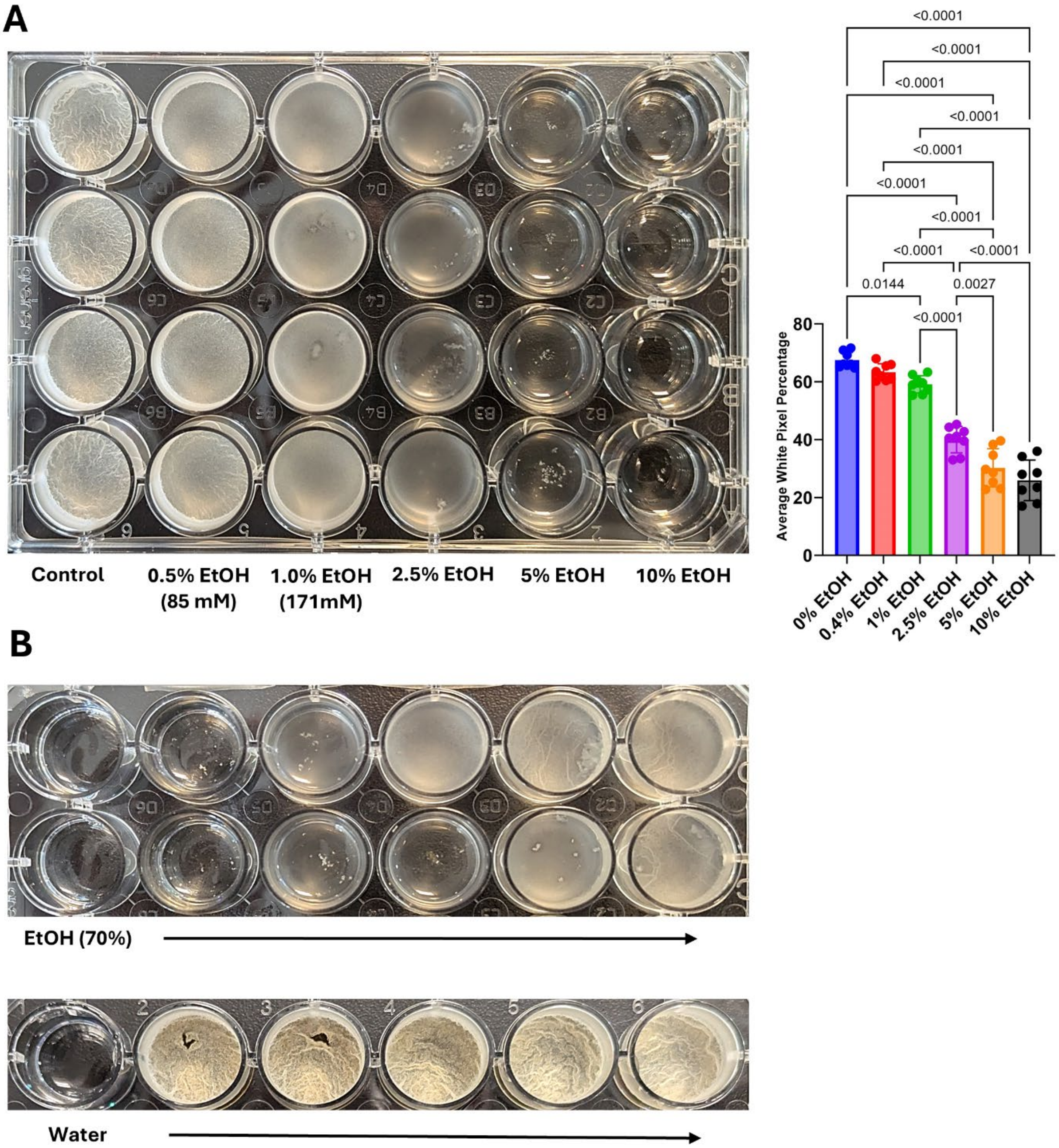


**Supplementary Figure 1: Ethanol modulates *B. subtilis* pellicle formation.**

**A**. Top-down view of *B. subtilis* pellicles formed in 24-well plates containing MSgg medium supplemented with increasing concentrations of ethanol (0–10% v/v) after ~24 h incubation at 30 °C. The corresponding quantification of pellicle surface coverage (right) is expressed as the average percentage of white pixels. Statistical significance was determined using one-way ANOVA (*P* values indicated). B. Comparison of pellicle formation under 70% ethanol solvent controls versus water controls, demonstrating the specific effect of ethanol concentration gradients on biofilm structural integrity.

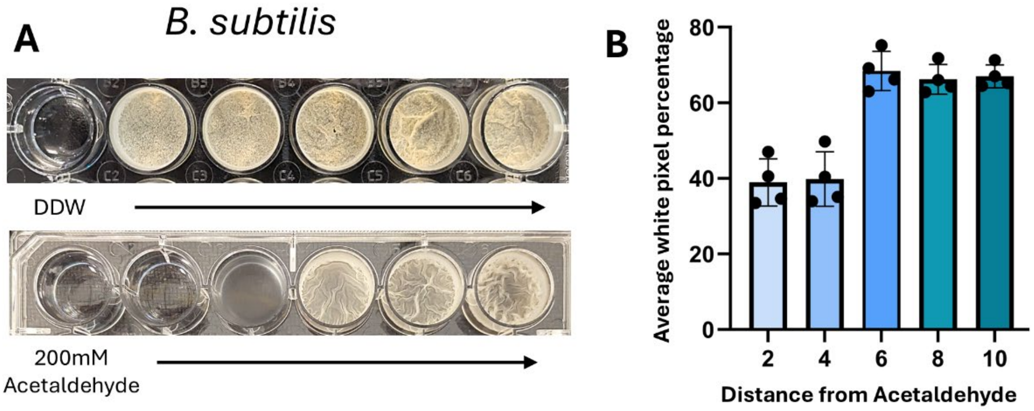


**Supplementary Figure 2: Effect of volatile acetaldehyde on biofilm formation**. (A) *Bacillus subtilis* pellicle formation in MSgg medium exposed to double-distilled water (DDW) as a control (top) and 200 mM acetaldehyde (bottom). (B) Quantification of *B. subtilis* pellicle robustness based on average white pixel percentage relative to the distance from the acetaldehyde source. Pellicle formation is significantly inhibited near the acetaldehyde wells.

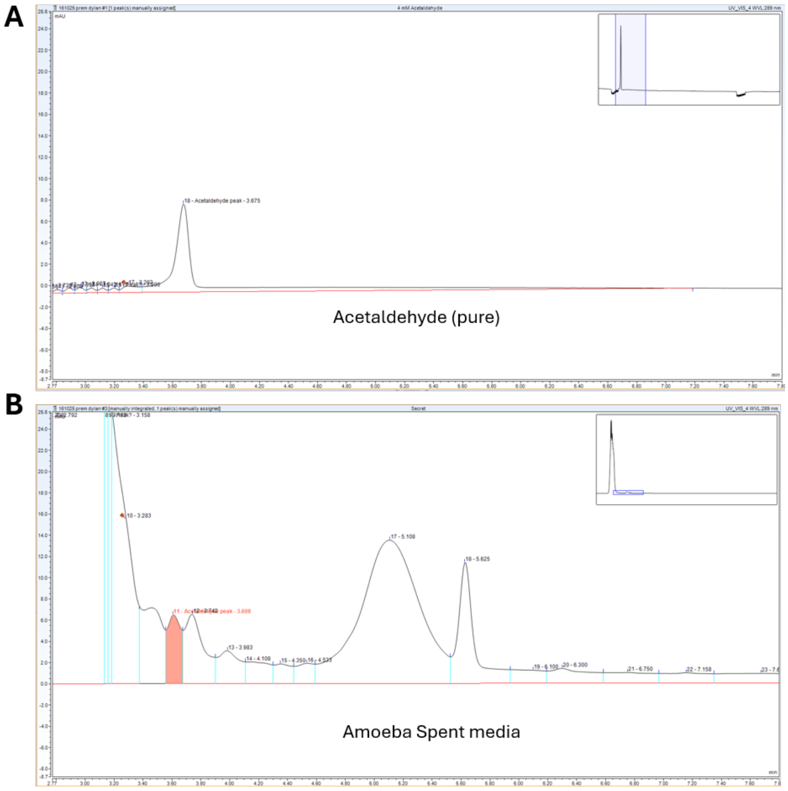


**Supplementary Figure 3:** Chromatographic profile showing a specific peak at the retention time of acetaldehyde.

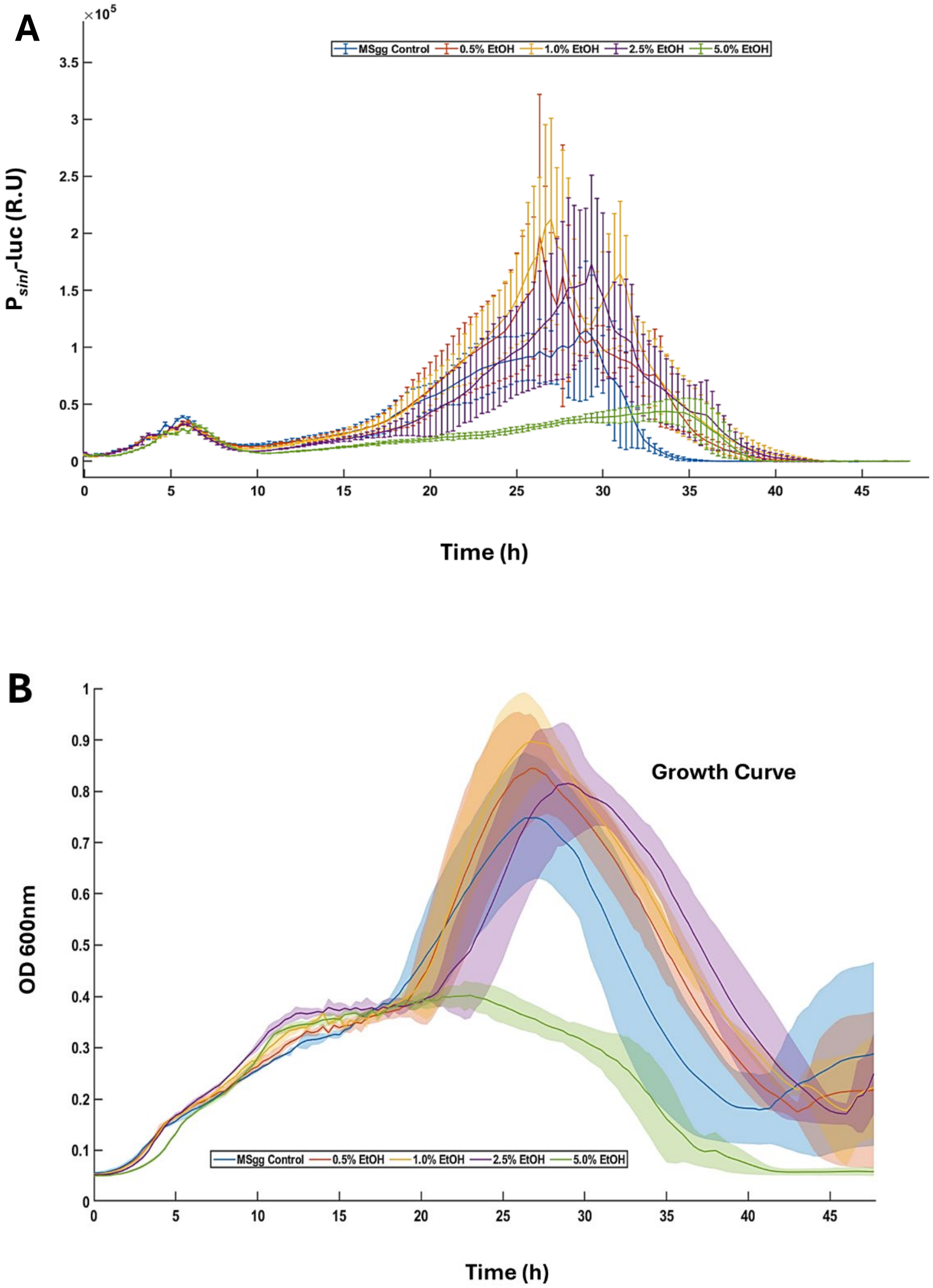


**Supplementary Figure 4:** Ethanol modulates *sinI* transcription. A. Time-dependent expression profiles of the biofilm master regulator reporter strain $P_{sinI}$-luc cultured in MSgg medium supplemented with the indicated concentrations of ethanol. Luminescence (relative units, R.U.) was monitored every 20 min. Error bars represent the SD (n=4). Decreased peak expression at 5.0% ethanol compared to the control. B. Growth curves (OD 600) of *B. subtilis* under the corresponding conditions. Shaded regions indicate SD (n=4). Ethanol extends the lag phase and reduces final biomass in a dose-dependent manner.

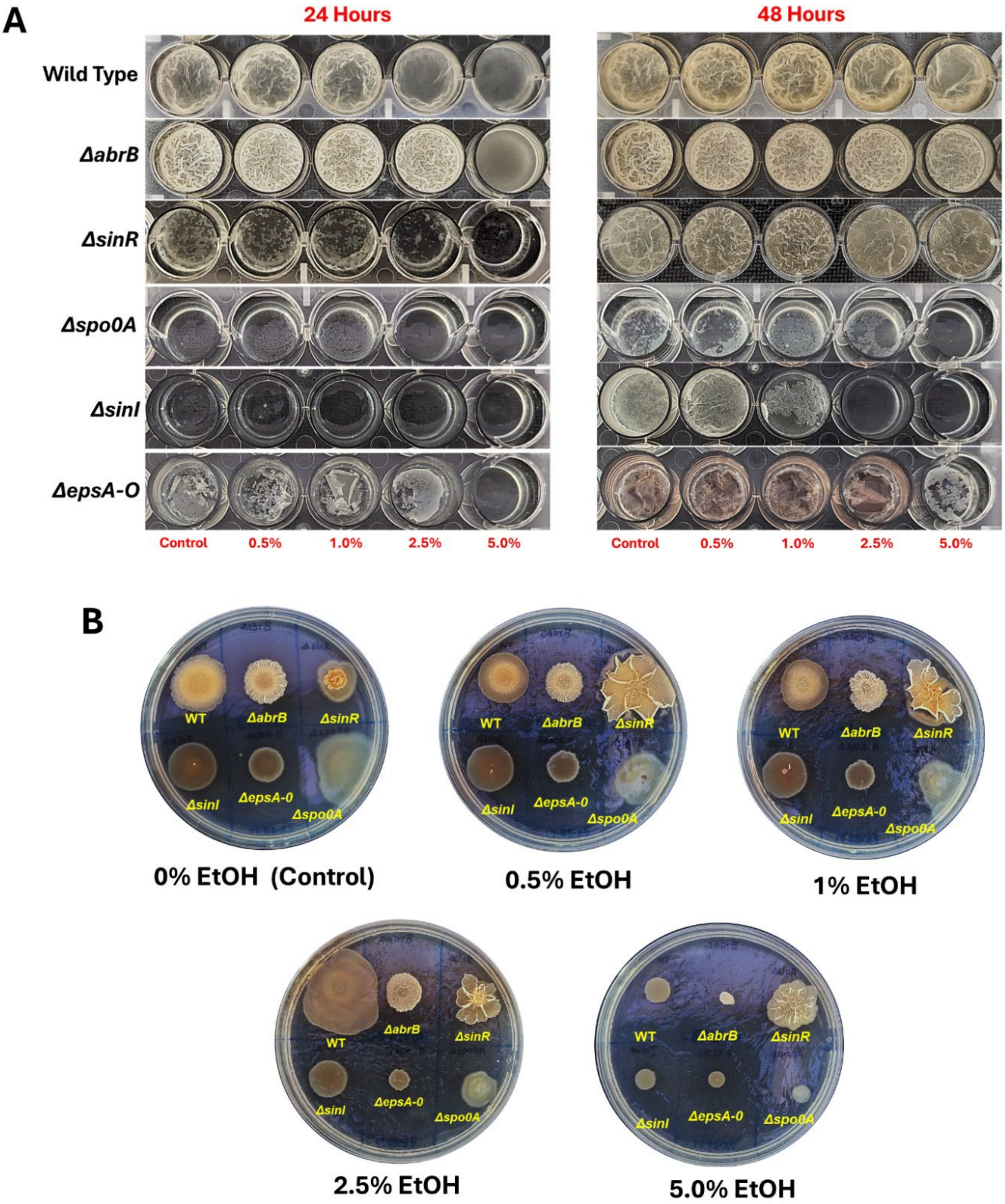


**Supplementary Figure 5: Ethanol inhibits biofilm formation.** A. Pellicle morphology of wild-type (WT) and biofilm regulator mutant strains (*ΔabrB, ΔsinR, Δspo0A, ΔsinI*, and matrix-deficient mutant *Δeps* cultured in MSgg medium with increasing ethanol concentrations (0–5%) at 24 h and 48 h. The hyper-wrinkled phenotype of *ΔsinR* persists, while upstream biofilm-regulatory mutants (*Δspo0A*) fail to form pellicles regardless of ethanol presence. B. Colony morphology of the indicated strains on MSgg agar plates supplemented with 0–5% ethanol, showing differences in colony architecture under ethanol stress.

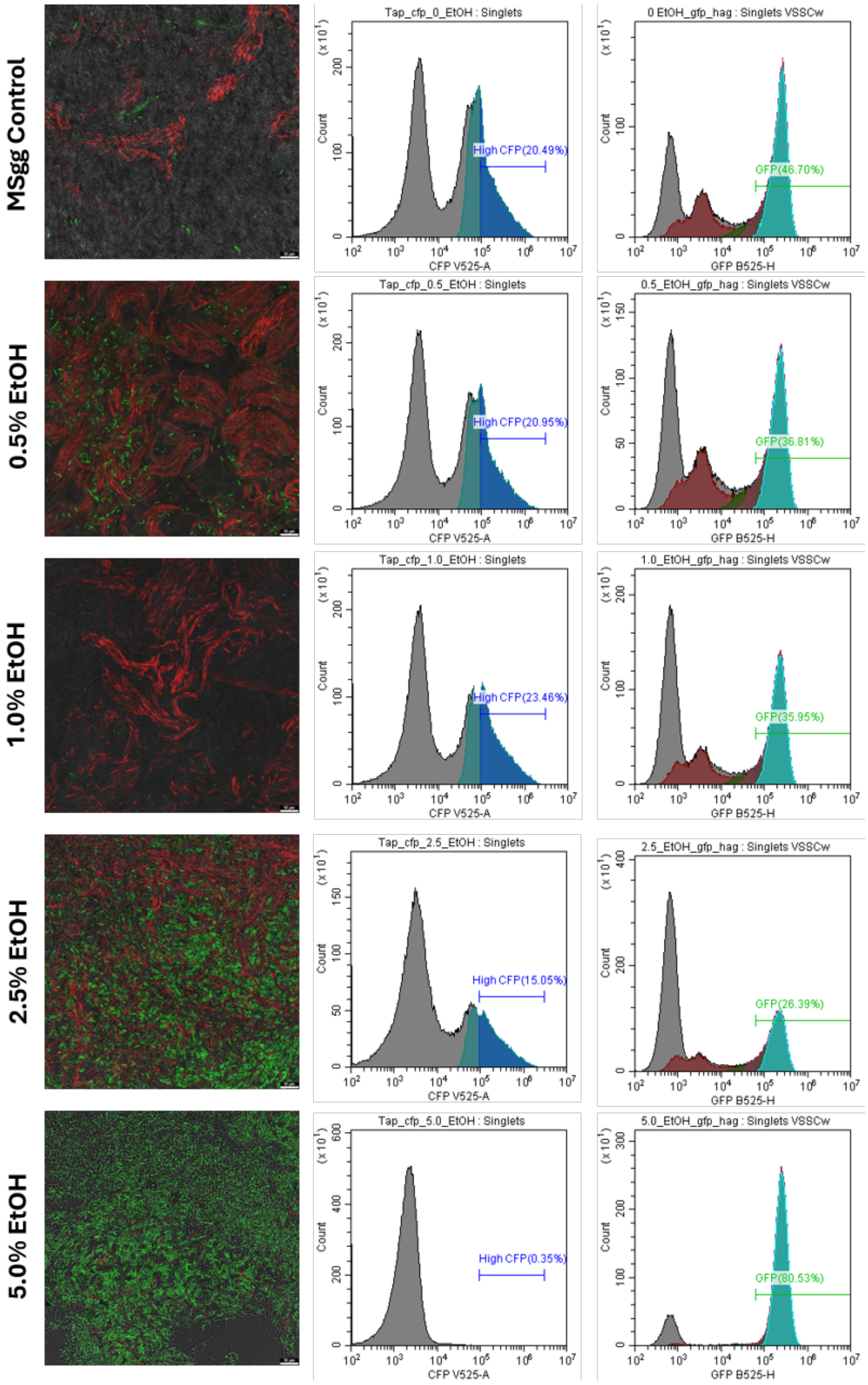


**Supplementary Figure 6**: **Ethanol-induced shifts in matrix and motility gene expression.** Confocal microscopy images showing the distribution of matrix-producing cells (TasA mCherry) and motile cells ($P_{hag}$-GFP) in control versus Ethanol-treated conditions. Flow cytometry histograms of $P_{tapA}$-CFP (matrix gene) and $P_{hag}$-GFP (flagellar gene) expression in MSgg control medium and under treatment with 0.5% to 5% EtOH. Percentages indicate the fraction of the population positive for the respective fluorescent reporters.

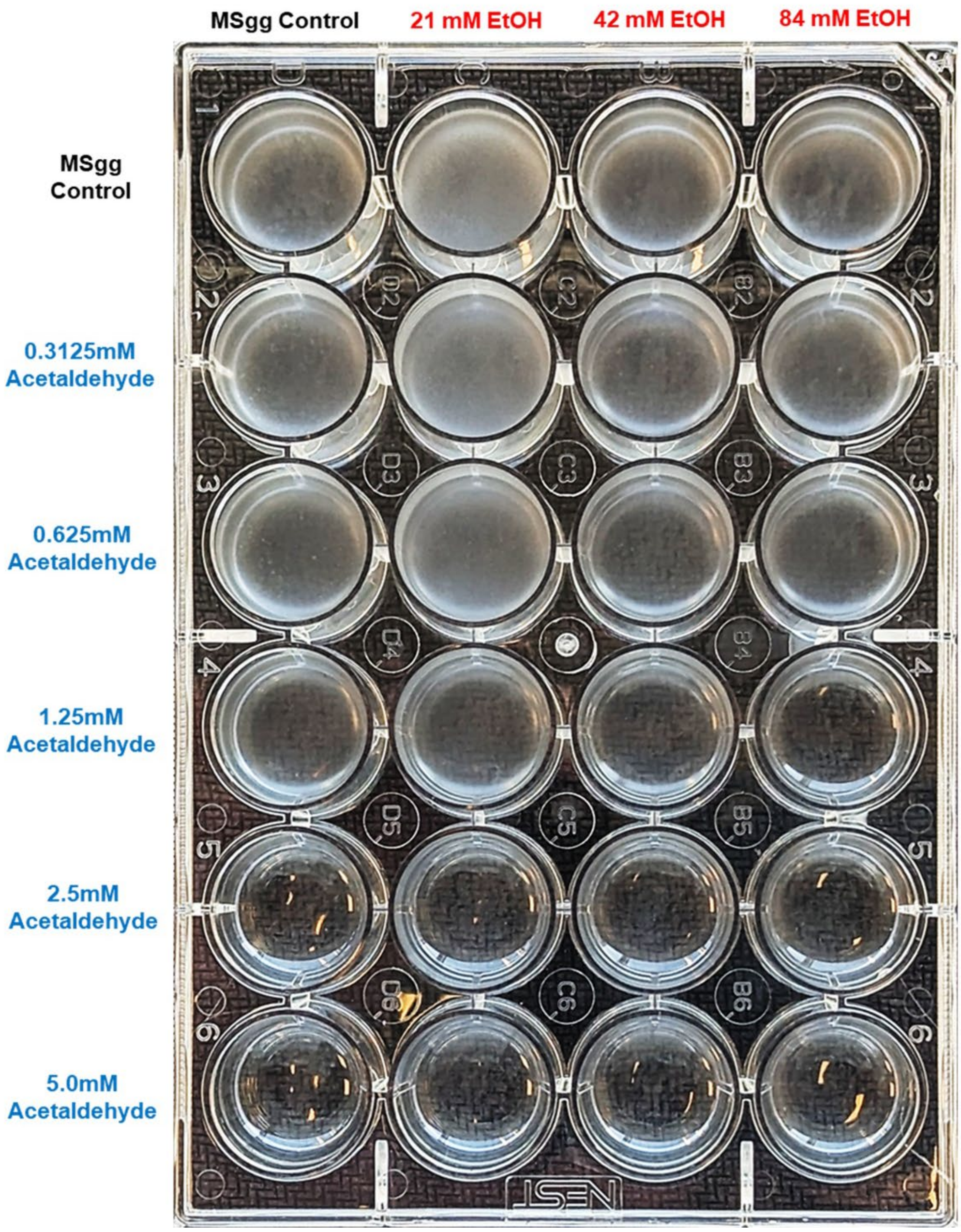


**Supplementary Figure 7: Combinatorial effects of acetaldehyde and ethanol on pellicle biofilm formation.** *B. subtilis* pellicles cultured for 24 h in MSgg medium arranged in a cross-gradient matrix of ethanol (columns, 0–84 mM) and acetaldehyde (rows, 0–5.0 mM). The addition of acetaldehyde to low ethanol concentrations progressively impairs biofilm development.

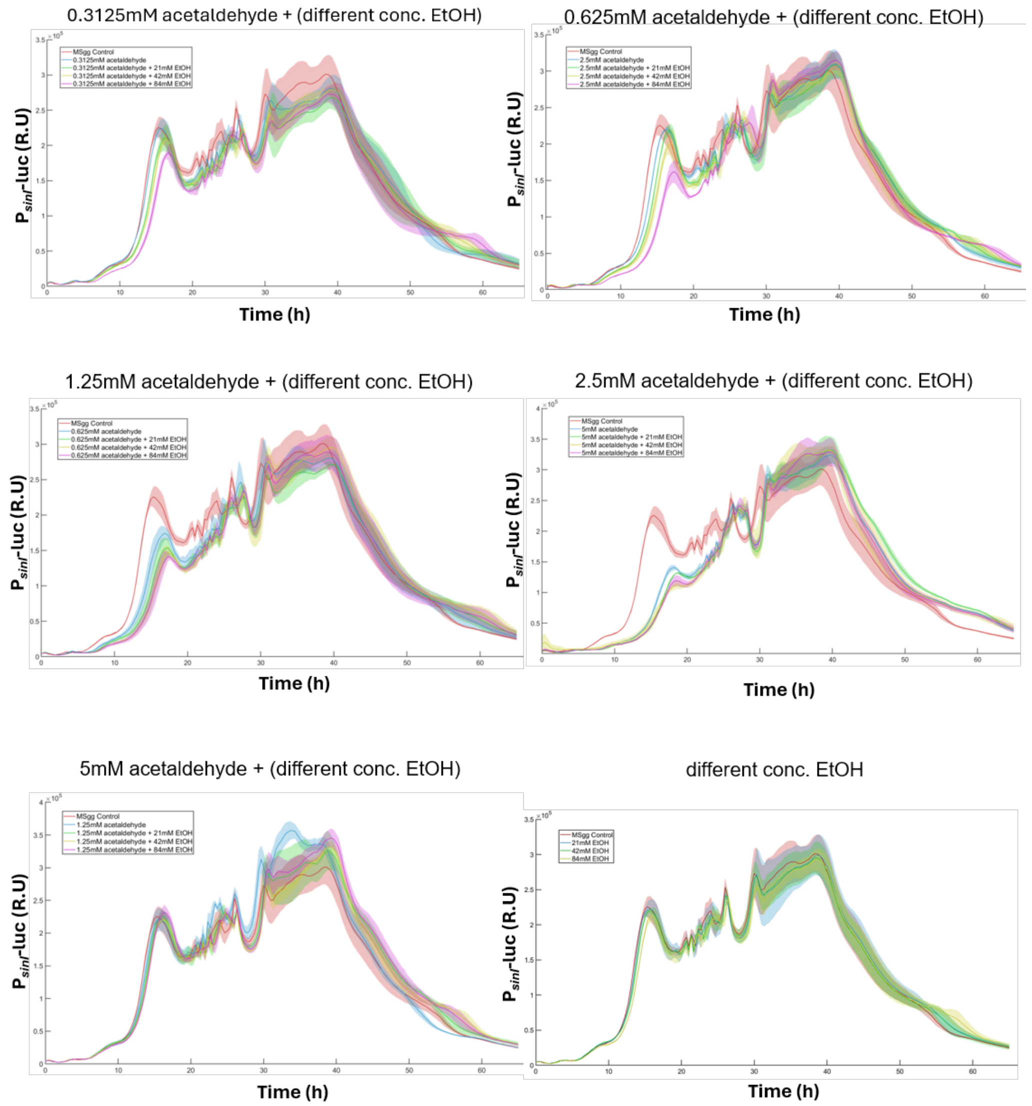


**Supplementary Figure 8: Combinatorial effects of acetaldehyde and ethanol on biofilm gene expression.** Luciferase activity of $P_{sinI}$-luc was measured in the presence of increasing concentrations of acetaldehyde (0.3125 mM to 5 mM) combined with varying concentrations of ethanol (0.5–5%). Titration of ethanol in the presence of acetaldehyde demonstrates that the metabolites synergistically enhance the inhibition of the *sinI* promoter expression.

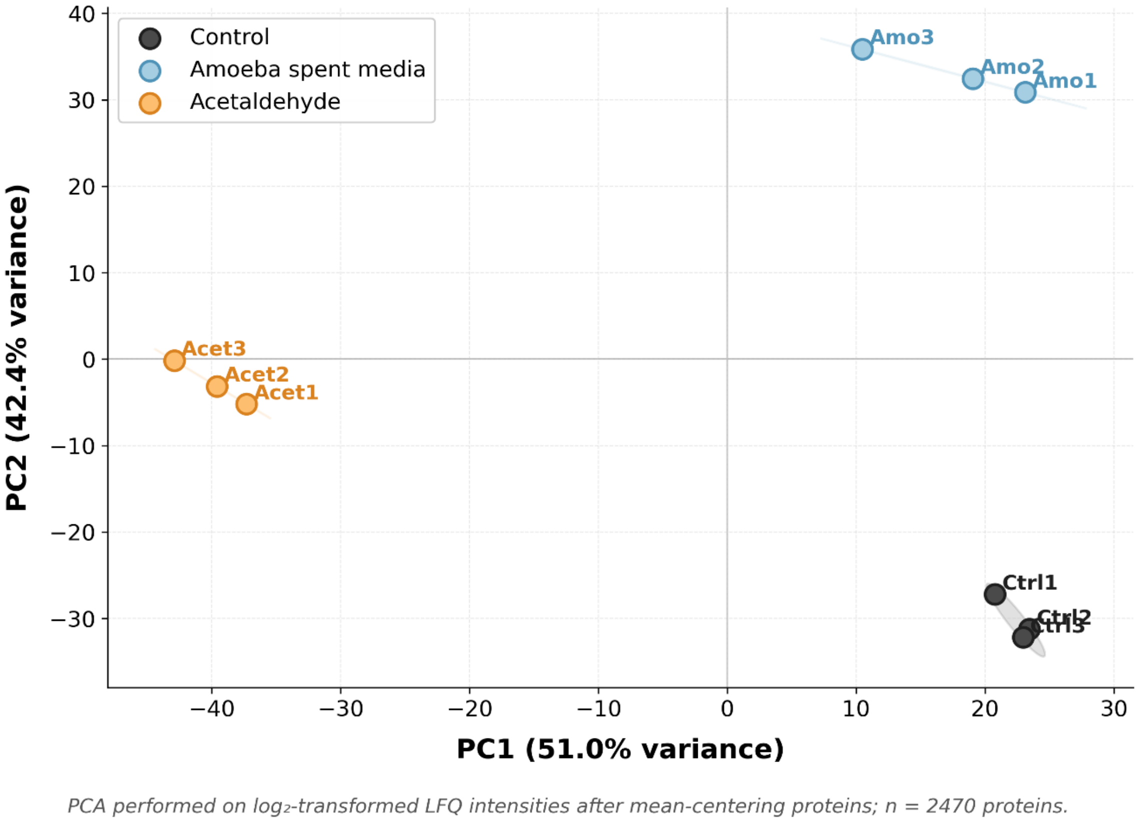


**Supplementary Figure 9: Principal component analysis (PCA) of label-free quantification (LFQ) proteomics**. Profiles from Control, Amoeba spent media, and Acetaldehyde-treated *B. subtilis*. Each point represents an individual replicate. PC1 explains 51% of the total variance and separates Acetaldehyde-treated samples from Control and Amoeba spent media samples, whereas PC2 explains 42.4% of the variance and separates Amoeba spent media from Control.

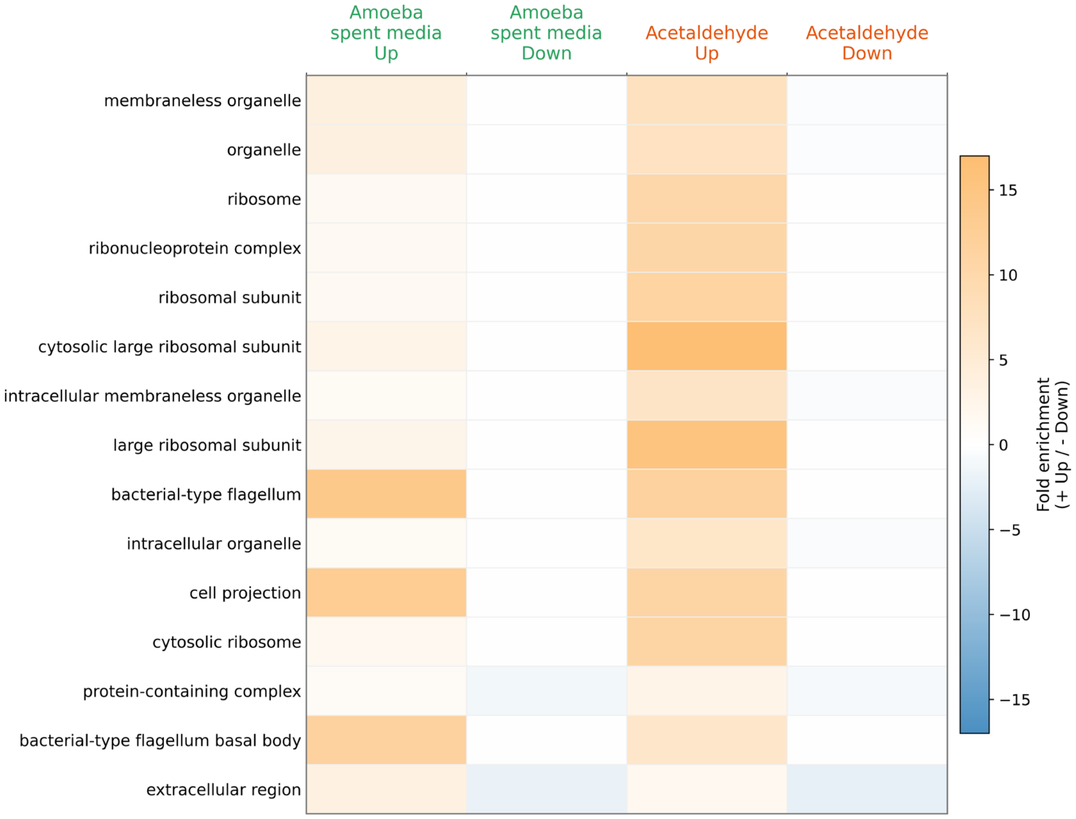


**Supplementary Figure 10: Gene Ontology cellular component enrichment matrix comparing Amoeba spent media and Acetaldehyde responses.** Columns represent upregulated and downregulated protein sets for each treatment. Orange indicates enrichment among upregulated proteins, blue indicates enrichment among downregulated proteins, and color intensity reflects enrichment magnitude

| **Strain** | **Reference** |
|---|---|
| *B. subtilis* NCIB 3610 | Branda *et al.,* 2001 [1] |
| *B. subtilis 168* | Mcloon et al., 2012[2] |
| *B. subtilis* lacking pBS32 | Rosenberg et al., 2016[3] |
| *ΔepsH*::*tet* | Oppenheimer-Shaanan *et al.,* 2016 [4] |
| *ΔsinR*::*spec* | Steinberg *et al.,* 2020 [5] |
| *ΔabrB::kan* | Bloom-Ackermann *et al.,* 2016 [6] |
| *Δspo0A*::*kan* | Bloom-Ackermann *et al.,* 2016 [6] |
| *ΔepsA-O*::*kan* | Steinberg *et al.,* 2020[5] |
| *ΔsinI*::*spec* | Bloom-Ackermann et al., 2016 [6] |
| *amyE*::$P_{sinI}$ *-lux* (Kan) | Oppenheimer-Shaanan et al., 2016[4] |
| *amyE*::$P_{hag}$ *-gfp* (cam) | Steinberg et al., 2020 [5] |
| *amyE*::$P_{tapA}$ *-cfp* (cam) | Steinberg et al., 2020 [5] |
| *amyE*::$P_{tapA-tapA-sipW-tasA-}$TasAmCherry (cam) | Steinberg et al., 2020 [5] |
| *B. subtilis* natto | Maan et al., 2022[7] |
| *Bacillus velezensis* FZB42 | Maan et al., 2022[7] |
| *B. megaterium isolate VS1* | Maan et al., 2022[3] |
| *B. subtilis PY79* | Maan et al., 2022[3] |
| *ΔcdaA* | Gilhar et al., 2024[8] |
| *ΔcdaS* | Gilhar et al., 2024[8] |
| *ΔcdaA ΔcdaS* | Gilhar et al., 2024[8] |
